# HydRA: Deep-learning models for predicting RNA-binding capacity from protein interaction association context and protein sequence

**DOI:** 10.1101/2022.12.23.521837

**Authors:** Wenhao Jin, Kristopher W. Brannan, Katannya Kapeli, Samuel S. Park, Hui Qing Tan, Maya L. Gosztyla, Mayuresh Mujumdar, Joshua Ahdout, Bryce Henroid, Katherine Rothamel, Joy S. Xiang, Limsoon Wong, Gene W. Yeo

## Abstract

RNA-binding proteins (RBPs) control RNA metabolism to orchestrate gene expression, and dysfunctional RBPs underlie many human diseases. Proteome-wide discovery efforts predict thousands of novel RBPs, many of which lack canonical RNA-binding domains. Here, we present a hybrid ensemble RBP classifier (HydRA) that leverages information from both intermolecular protein interactions and internal protein sequence patterns to predict RNA-binding capacity with unparalleled specificity and sensitivity using support vector machine, convolutional neural networks and transformer-based protein language models. HydRA enables Occlusion Mapping to robustly detect known RNA-binding domains and to predict hundreds of uncharacterized RNA-binding domains. Enhanced CLIP validation for a diverse collection of RBP candidates reveals genome-wide targets and confirms RNA-binding activity for HydRA-predicted domains. The HydRA computational framework accelerates construction of a comprehensive RBP catalogue and expands the set of known RNA-binding protein domains.

**Highlights:**

- HydRA combines protein-protein interaction and amino acid sequence information to predict RNA binding activity for 1,487 candidate genes.
- HydRA predicts RNA binding with higher specificity and sensitivity than current approaches, notably for RBPs without well-defined RNA-binding domains.
- Occlusion Mapping with HydRA enables RNA-binding domain discovery.
- Enhanced CLIP confirms HydRA RBP predictions with RNA-binding domain resolution.

## INTRODUCTION

Individually or in ribonucleoparticle protein (RNP) complexes, RNA binding proteins (RBPs) orchestrate every step of the RNA life cycle ^1^. The recognition that an increasing number of human diseases are associated with perturbed RBP function ^2–4^ has fueled a resurgence in RBP discovery efforts. In the past decade, RNA-interactome capture and organic phase separation-based technologies have uncovered thousands of proteins that contact RNA^5,6^. These experimental RBP-discovery approaches reveal that the RNA-binding proteome is cell type- and context-specific, indicating that completing the catalog of known RBPs will require resource-intensive RNA-interactome interrogations across many tissues and conditions. Furthermore, approximately half of the recently discovered RNA-interacting proteins are unconventional RBPs that lack both annotated RNA-related biological functions and annotated RNA-binding domains ^5,6^, which poses problems for prioritizing candidates for validation and characterization.

An exciting strategy to these affinity-based and phase-separation-based RNA-interactome capture assays is to utilize machine or deep learning approaches to predict proteome-wide RNA-binding capacity. RBP classifiers that utilize primary sequence information, known as sequence-based classifiers, such as RNApred ^7^, SPOT-seq ^8^, catRAPID signature ^9^, RBPPred ^10^ and TriPepSVM ^11^ have been developed that require only sequence information as input. Apart from SPOT-seq, all of these classifiers utilize classical machine learning models, such as Support Vector Machines (SVMs), trained on features extracted from protein sequence such as amino acid composition, *k*-mers (the set of all the *k*-length sequences) and predicted physicochemical properties to distinguish RBPs from non-RBPs. Each of these algorithms suffers from its own notable limitations. Machine learning based methods that rely on human-defined features such as global protein properties (e.g. amino acid compositions) and primary sub-sequence count (e.g. *k-mer*) are not sensitive to relatively small regions of RNA binding. SPOT-seq, instead, uses a template-based strategy that aligns query proteins to known protein-RNA complex structures and evaluates query protein RNA-binding affinity ^8^. However, SPOT-seq depends on the quality and comprehensiveness of the template set of protein-RNA complex structures. While these classifiers categorize “conventional” RBPs (with known RNA-binding domains) with some degree of accuracy, they demonstrate limited predictive ability for the entire catalog of experimentally defined human RBPs, of which > 50% are anticipated to be unconventional RBPs lacking known RNA-binding domains ^11^. This limited predictive ability is due to the simplicity of the sequence-based machine learning models employed by these classifiers as well as the inherent complexities of RBP prediction. In practice, low numbers of positive training samples prevent the modeling of the implicit and complicated sequence patterns that exist in unconventional RBPs.

To identify unconventional RBPs, we previously developed a machine learning algorithm termed SVM Obtained from Neighborhood Associated RBPs (SONAR) that utilized protein-protein interaction (PPI) networks ^12^. SONAR leveraged our empirical findings that RBPs often physically interact with one another (either in RNA-dependent or –independent mode) and identified unconventional RBPs with a high validation rate. However, SONAR failed to classify a portion of conventional RBPs with known RNA-binding domains, likely because SONAR’s high dependency on experimentally defined PPI networks limits its application for proteins underrepresented in these networks. Here we set out to develop computational algorithms capable of recognizing both conventional and unconventional patterns exclusive to RBPs by integrating both external (PPI-based) and internal (sequence-based) information layers.

State-of-the-art machine learning algorithms, such as deep learning, have not only improved the precision of predictive models but have revolutionized the way data is processed. For example, convolutional neural network (CNN), a widely used class of deep learning algorithms in computer vision (CV), and Transformer, powerful for natural language processing (NLP) problems, are able to extract useful information automatically from raw data such as photographs or text and make predictions without hand-engineered and intrinsically biased features. Several applications have been developed with CNN and Transformer to solve biological problems such as protein folding^13^, SNP calling^14^, and protein family classification ^13^, demonstrating their predictive strengths in biological sequence classification. However, to avoid overfitting and maintain robustness of these models, large training datasets are necessary. To alleviate this problem of small positive training sizes, pretraining methods such as self-supervised learning and transfer learning have been used successfully for CV and NLP implementations ^14^. These methods have also been applied in protein classification problems. A Transformer-based protein language model named ProteinBERT recently achieved state-of-the-art performance on multiple protein classification tasks covering diverse protein properties (including protein structure, post translational modifications and biophysical attributes) ^15^. Traditional machine learning models, such as SVM and Random Forest, work well on small datasets and address the problem of overfitting. Thus, intuitively, ensemble learning that integrates deep learning with traditional machine learning models offers a promising strategy to design more precise and robust RNA-binding classifiers.

Here, we introduce a new machine learning-based algorithm, **Hy**bri**d** ensemble classifier for **R**N**A**-binding proteins (HydRA), that predicts not only the RNA-binding capacity of proteins but also regions of proteins that are involved in RNA-protein interaction. This algorithm applies an ensemble learning method that integrates convolutional neural network (CNN), Transformer and SVM in RBP prediction by utilizing both intermolecular protein context and sequence-level information. Pretraining methods including self-supervised and transfer learning are used to alleviate model overfitting and increase model robustness in the CNN and Transformer. We show HydRA outperforms all available state-of-the-art RBP classifiers and additionally recognizes known RNA-binding related elements with Occlusion Map, a model interpretation approach originally developed in the computer vision arena. Our results also suggest that intermolecular protein context is critical for identifying novel RBPs whose sequence or structural patterns may not be easily captured.

In summary, HydRA predicted 1,487 unclassified RBPs. Using enhanced CLIP (eCLIP), we validated a subset of HydRA predictions, including previously reported but not characterized RBPs HSP90A and the YWHA family of proteins, as well as the novel RBP candidates INO80B, PIAS4, NR5A1, ACTN3, and MCCC1, corroborating bona-fide RBPs and providing molecular insights into biological functions. For these RBP candidates, we experimentally verified RNA-binding activity for novel RNA-binding domains predicted by Occlusion Map. These results showcase HydRA as the highest accuracy RBP classifier to date, and the first capable of predicting RNA-binding domains from unconventional RBP candidates at amino-acid resolution.

## RESULTS

### Improved SONAR models penalize low PPI neighborhood and increasing network density

We previously developed SONAR (1.0), an SVM-based machine learning algorithm that leverages large-scale experimental protein-protein interaction (PPI) data (without protein sequence information) to predict unannotated RBPs ^16^. Employing the BioPlex PPI dataset ^17^, SONAR was able to predict RBPs with high specificity and sensitivity with BioPlex proteins and previously compiled RBP list. The success of SONAR was based on observations that an unusually high proportion of proteins that associate with RBPs (via RNA-dependent and independent interactions) are themselves RBPs. As such, SONAR’s ability to accurately identify RBPs depended on the density of protein-protein interactions centered on an interrogated protein (the number of “edges”) or the completeness of the local PPI network. To illustrate, known RBPs that were assigned low SONAR scores (false negatives) harbored significantly fewer PPI edges than correctly predicted known RBPs (true positives) (**Figure 1A**). To evaluate the extent that the density of PPI networks affects the predictive power of SONAR, we constructed a more comprehensive PPI network by combining BioPlex2.0 ^17^ with Mentha ^18^, a collection of physical protein-protein interaction from MINT ^19^, BioGRID ^20^ and IntAct ^21^. We refer to this ensemble network as Mentha-BioPlex (MB). Next, we incrementally reduced the network density of Mentha-BioPlex by randomly removing a percentage of edges and determined the ROC-AUC value of SONAR by 10-fold cross-validation. Indeed, both sensitivity and specificity (measured by ROC-AUC values) of SONAR improves with more comprehensive PPI networks (**Figure 1B**). To evaluate the predictive power of SONAR on a corrupted network, we incrementally removed edges from a shuffled Mentha-BioPlex network such that the number of edges were unchanged, but edges were randomly connected to maintain network topological properties (i.e. node degree distribution). The resulting ROC-AUC values of SONAR models using shuffled Mentha-BioPlex networks were significantly lower (< 0.65) with large standard error of the mean compared with that of the unshuffled network (**Figure 1B**). This result demonstrates that low quality PPI data will weaken the predictive power of SONAR. To alleviate the impact of PPI sparsity on SONAR performance, we added an indicator feature in our algorithm to mark proteins with limited PPI information, which led to an increase in ROC-AUC value by 0.016 (from 0.767 to 0.783) for the SONAR model with BioPlex network (**Figures 1C** and **1D**) and an increase in Precision-Recall-Area Under Curve (PR-AUC) value by 0.108 (from 0.347 to 0.455) (**Figure S1B**). We refer to this revised model as SONAR2.0. By replacing BioPlex network with the denser Mentha-BioPlex network (**Figures 1C** and **S1A**), we improve SONAR1.0’s performance by another 0.078 (from 0.783 to 0.861, **Figure 1D**) in ROC-AUC and 0.108 (from 0.455 to 0.563) in PR-AUC (**Figure S1B**).

**Figure 1.**
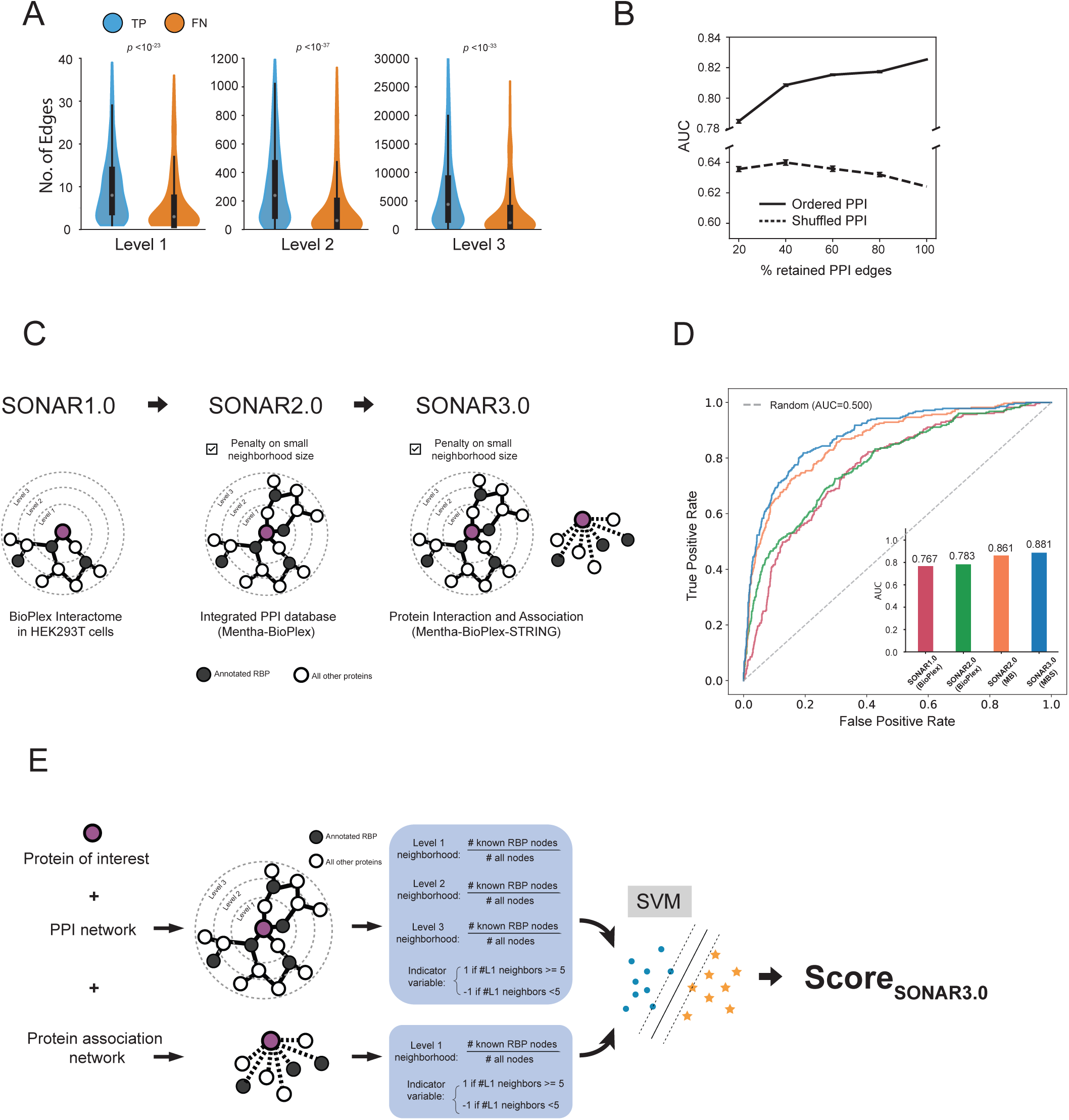
Protein Interaction and Association based (SONAR3.0) classifier presents upgraded RBP classification performance from SONAR. **A.** Violin plots showing the number of Level 1, Level 2, and Level 3 neighbors for RBPs that were scored correctly (true positive) or incorrectly (false negative) by SONAR. The median and mean between true positive and false negative RBPs is significant for all levels (Level 1, p<1e-23; Level 2, p<1e-37; Level 3, p<1e-33 by one-side Mann-Whitney U test). The median of each population is shown. **B.** Positive correlation between network density and ROC-AUC value. Random shuffling of edges resulted in no correlation with ROC-AUC value. Error bars represent variability from 10 replicates. **C.** The evolution from SONAR1.0 to SONAR3.0 where new PPI data type was introduced in each version. **D.** Schematic of the PIA-based classifier that takes features extracted from protein-protein interaction (PPI) network and protein association network as input. Support vector machine (SVM) with radial basis function (RBF) kernel was employed for mathematical modelling. The local PPI network (the first three levels of neighborhoods) and the local protein association network (only the direct neighbors) are first retrieved for the protein of interest. Features are obtained by calculating the percentage of known RBPs in each neighborhood and an indicator variable is added for each network to penalize the case with low number of protein neighbors. **E.** ROC-AUC analysis to compare the predictive power of SONAR1.0 with BioPlex network, SONAR2.0 with BioPlex network, SONAR2.0 with mentha-BioPlex (MB) network and SONAR3.0 with mentha-BioPlex and STRING (MBS). Inset: Bar graph of ROC-AUC values

Next, we evaluated if adding functional protein association data from the *Search Tool for the Retrieval of Interacting Genes/Proteins* (STRING) database would further improve SONAR2.0 as functional protein association data had previously been successfully applied in other PPI-based protein prediction tasks^22^. Like SONAR2.0, we set an indicator feature to penalize proteins with few interacting neighbors within its local neighborhood in the STRING network. With the extended protein networks, we obtained an improved RBP classifier (SONAR3.0) using an SVM model with a radial basis function (RBF) kernel (**Figures 1E** and **S1A**), where the prediction performance increased by 0.020 in ROC-AUC and 0.051 in PR-AUC (**Figures 1D** and **S1B**). Specifically, the precision-recall analysis highlights the advantage of incorporating more protein-protein interaction and functional association information to densify our networks and minimize false positives (**Figure S1B**). For instance, the precision of the model increased from 0.41 to 0.57, 0.73 and 0.76 at the 0.3 recall level. Setting a cutoff to recover the top 30% of known RBPs ranked by classification scores, we obtain 86 false positive predictions (precision = 0.41) using SONAR1.0, 43 false positives using SONAR1.0 with an updated BioPlex network, 21 false positives using SONAR2.0 (with Mentha-BioPlex) and only 18 false positives using SONAR3.0 (**Figure S1B**).

### HydRA-seq classifies RNA-binding proteins from protein sequence by ensemble learning

We next set out to augment the HydRA workflow to enable classification of RBPs and non-RBPs based on the combination of two components: 1) SONAR3.0 which captures the RNA-binding-related intermolecular patterns (**Figure 1E**), and 2) RNA-binding-related patterns from primary amino acid sequences (referred to as HydRA-seq) (**Figures 2A-2D**). The primary amino acid sequence of protein domains has been previously leveraged by RBP classifiers with classic machine learning models to predict RNA binding capacity of canonical RBPs with well-defined RNA binding domains ^7–11^. Expectedly, these sequence-based classifiers perform poorly on experimentally defined RBPs that do not have canonical RNA binding domains (**Figures S2A** and **S2B**).

**Figure 2.**
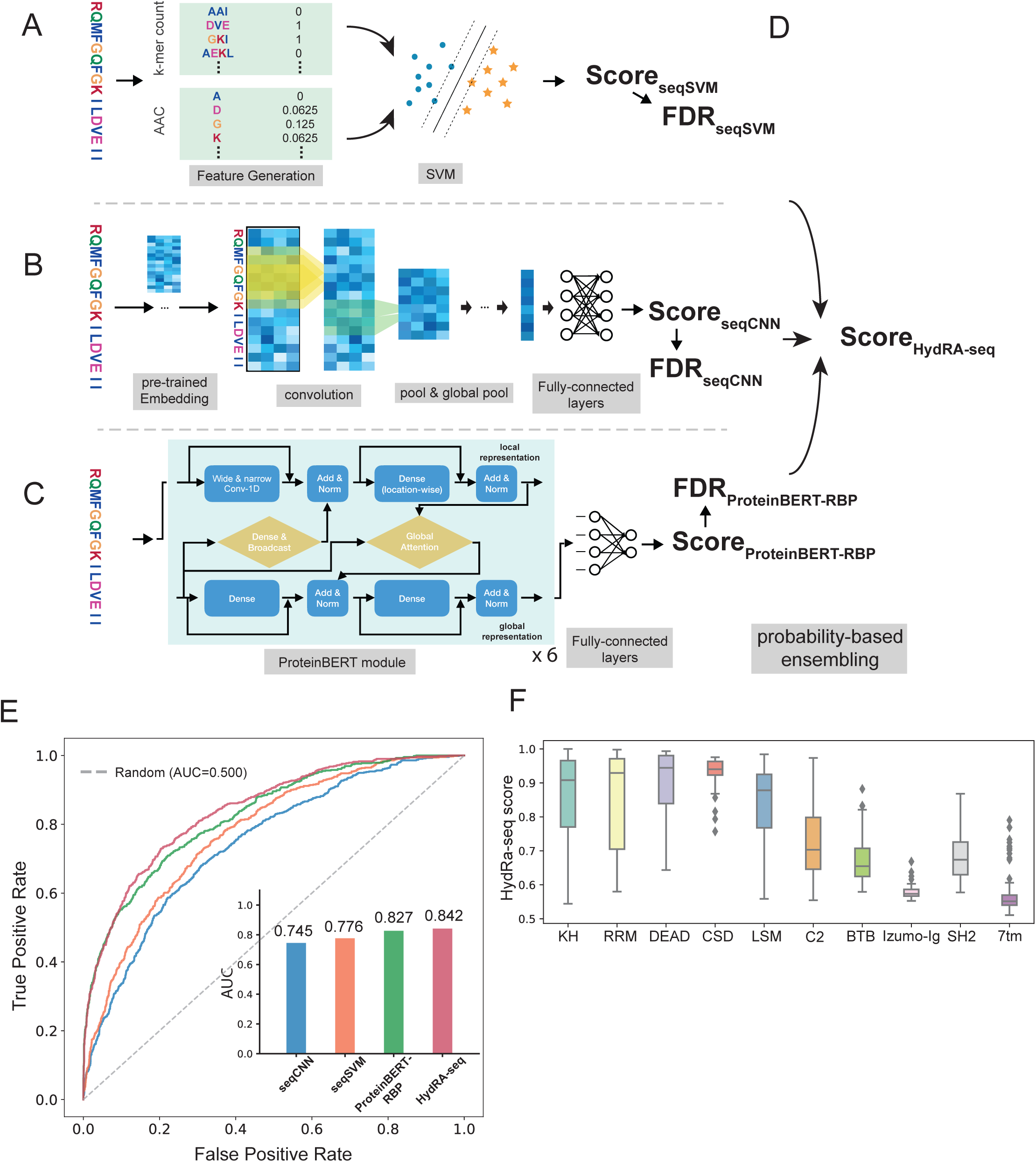
HydRA-seq classifies RNA-binding proteins. **A-D.** HydRA-seq model that consists of sub-classifiers including seqSVM, seqCNN and ProteinBERT. **A.** Schematic of seqSVM model that is based on SVM with RBF kernel employing k-mer, predicted secondary structures and amino acid composition in the prediction. **B.** Schematic of seqCNN model that is comprised of 1 embedding layer, 2 set of convolutional and pooling layers, 1 global pooling layer, 2 fully connected layers and 1 output layer and takes protein sequence and predicted secondary structures as input. Only 1 set of convolutional and pooling layers are shown. Each layer (except embedding and pooling layers) is coupled with a batch-normalization layer which is not shown in the diagram. **C.** Schematic of ProteinBERT-RBP classifier. The query protein sequence was first fed into 6 sequential ProteinBERT modules in each of which two parallel neural networks were applied to extract and process the sequence (i.e. local presentation) and functional properties (i.e. global presentation) of the protein respectively while attention mechanism were used to integrate these two types of information. The output embedding vector was then conveyed into a fully connected neural layer to generate the classification score (Score_ProteinBERT-RBP_). **D.** Probability-based ensembling approach that convert the classification scores from the three RBP classifiers into the overall sequence-based RBP score (Score_HydRA-seq_). False discovery rate of each classifier’s prediction was calculated for each protein and was used to generate the Score_HydRA-seq_. **E.** We used ROC-AUC analysis to evaluate the performance of seqCNN, seqSVM, ProteinBERT and the ensemble classifier (denoted as “HydRA-seq”) of these three classifiers. Inset: Bar graph comparing ROC-AUC values. Inset: Bar graph comparing ROC-AUC values. **F.** Classification scores that are generated by the ensemble classifier of seqCNN, seqSVM and ProteinBERT for the amino acid sequences from five RNA-binding domains (RBDs) and five domains unrelated to RNA processing (non-RBDs). Th five RBDs are K homology (KH) domain, RNA recognition motif (RRM) domain, DEAD box (DEAD) domain, cold-shock domain (CSD) and like-SM (LSM) domain. The non-RBDs are C2 domain, BTB domain, Src homology 2 (SH2) domain, immunoglobulin (Ig) domainand 7 transmembrane (7tm) domain.

To combine the strength of both traditional machine learning models and deep learning models, we constructed three RBP classifiers with state-of-the-art machine learning technology using classical and novel sequence-based features which are generated from protein sequence. In the first classifier termed seqSVM, *k*-mer (sub-sequence of length *k*) and amino acid composition extracted from the amino acid sequences were used to train a support vector machine (SVM) with a radial basis function kernel (**Figure 2A**). For the second classifier called seqCNN, we employed a convolutional neural network (CNN) to learn features of RBPs from full-length amino acid sequences (**Figure 2B**). Briefly, an embedding layer pre-trained by BioVec ^23^ was employed to obtain a comprehensive representation of the biophysical and biochemical properties from each protein sequence, followed by convolutional and pooling layers that automatically extract and compress sequence-level features. Fully connected layers with non-linear activation units were added to further process the compressed features and generate final predictions. To avoid convergence of suboptimal solutions and accelerate the training process, seqCNN was initialized with weights from a convolutional autoencoder pretrained with all the protein sequences in our dataset before model training (see Methods). For the third classifier called ProteinBERT-RBP, protein sequences are fed into an attention-based Transformer architecture designed for proteins (ProteinBERT) ^15^ to obtain predicted RNA-binding propensities (**Figure 2C**). ProteinBERT-RBP was pretrained with ∼106 million UniRef90 protein sequences and their corresponding functional annotations from the Gene Ontology database in a self-supervised manner, and then fine-tuned using our human RBP annotations. As a result, ProteinBERT-RBP digests each input protein sequence in two parallel paths, (1) local features of the sequence and (2) global function of the sequence based on the knowledge obtained in the pretraining stage, while the two paths interact and guide the learning process of each other via attention mechanism and generate predictions on RNA-binding propensity.

ROC-AUC and precision-recall analysis with the held-out test set demonstrates good predictive power of seqCNN, seqSVM and ProteinBERT-RBP for individually identifying RBPs (**Figures 2E** and **S2C**). These classifiers identify RBPs with characterized RNA targets (CLIP-based data, collected in CLIPdb ^24^) or known RNA-binding or RNA-processing related domains (referred to as “characterized RBPs”) with approximate ROC-AUC values of 0.80, but performance is diminished for identifying RBPs lacking known RNA-binding domains (referred to as “uncharacterized RBPs”) (**Figures S2D** and **S2E**). To improve the prediction performance for both characterized and uncharacterized RBPs we designed a false discovery rate (FDR)-based ensemble approach to integrate the predictions from seqCNN, seqSVM and ProteinBERT-RBP (**Figure 2D**). This strategy utilizes the joint probability of each classifier’s predictions of being false discoveries and outputs the complementary probability of this joint probability as the ensemble classification score (more details in Methods). We referred to this ensemble classifier as HydRA-seq and find that HydRA-seq outperforms all of its sub-components (i.e. seqCNN, seqSVM and ProteinBERT-RBP) (ROC-AUC=0.842 and PR-AUC=0.643, **Figures 2E** and **S2C**). In particular, HydRA-seq exhibits better predictive power for identifying uncharacterized RBPs (ROC-AUC=0.782 and PR-AUC=0.347) when compared to individual seqCNN, seqSVM and ProteinBERT-RBP predictions (**Figures S2D** and **S2E**). HydRA-seq, trained with data only from the human proteome, also showed better performance on both characterized and uncharacterized RBPs than the current state-of-the-art sequence-based RBP classifier, TriPepSVM, which was trained with all the RBPs and non-RBPs in *human*, *Salmonella* and *E.coli* (**Figures S2D** and **S2E**, HydRA-seq: ROC-AUC = 0.913 and PR-AUC = 0.693 for characterized RBPs; TriPepSVM: ROC-AUC = 0.890 and PR-AUC = 0.508 for characterized RBPs, ROC-AUC = 0.719 and PR-AUC = 0.293 for uncharacterized RBPs).

We next tested whether HydRA-seq can distinguish known RNA-binding domains from other protein domains. We computed HydRA-seq classification scores for sequences collected from Pfam database corresponding to RNA binding domains (K homology domain, RNA recognition motif, DEAD box, cold-shock domain and like-SM domain) in different species. Expectedly, these sequences generally resulted in high classification scores (**Figure 2F**). In contrast, sequences from protein domains that are not commonly found in RBPs such as protein-protein interaction (BTB, IgG), membrane associated (transmembrane, C2), and src homology 2 domains overall had lower classification scores than the known RBDs (**Figure 2F**). These results demonstrate the capacity of our sequence-based RBP classifiers to distinguish sequence patterns that contribute to RNA-binding activity with domain level resolution.

### HydRA achieves state-of-the-art RBP classification performance

Given that SONAR3.0 and HydRA-seq individually predict RBPs with high accuracy even though they are trained using different protein features, we reasoned that a combined model would show enhanced performance compared to the individual ones. Indeed, we observed that combining prediction scores from SONAR3.0 and HydRA-seq components (i.e. seqCNN, seqSVM and ProteinBERT-RBP) using the FDR-based probabilistic ensemble approach (**Figure 3A**), produced a new model (herein termed HydRA) which outperformed its sub-classifiers with higher specificity and sensitivity displayed by AUC of 0.895 in ROC analysis and an AUC of 0.745 in precision-recall analysis within the hold-out test set (**Figures S3A** and **S3B, Table S1**). We next compared the performance of HydRA with other published RBP classifiers, namely SONAR, TriPepSVM, SPOT-seq, RNApred, catRAPID signature and RBPPred, and observed higher specificity and sensitivity than all other classifiers in the hold-out test set as shown in ROC-AUC and PR-AUC analyses (**Figures 3B** and **S3C**). High-quality protein structures predicted from amino acid sequences have recently become available for most natural proteins with AlphaFold2^25^. To test if predicted protein structures can further improve our RBP predictions, we constructed a group of structure-based RBP classifiers with AphalFold2 predictions as input using a graph neural network widely used in structure-based models for protein prediction problems ^26–28^. Optimal models (referred to as strucGNN1) were chosen after iterative feature selection and model tunning with our validation set (**Table S4**). We also obtained another structure-based RBP classifier by finetuning a pre-trained GNN-based protein model with our training set (see Methods), referred to as strucGNN-MIF. These strucGNNs achieve 0.813 - 0.827 in ROC-AUC analysis and 0.563 – 0.607 in PR-AUC analysis in the hold-out test set (**Figures 3B** and **S3C**).

**Figure 3.**
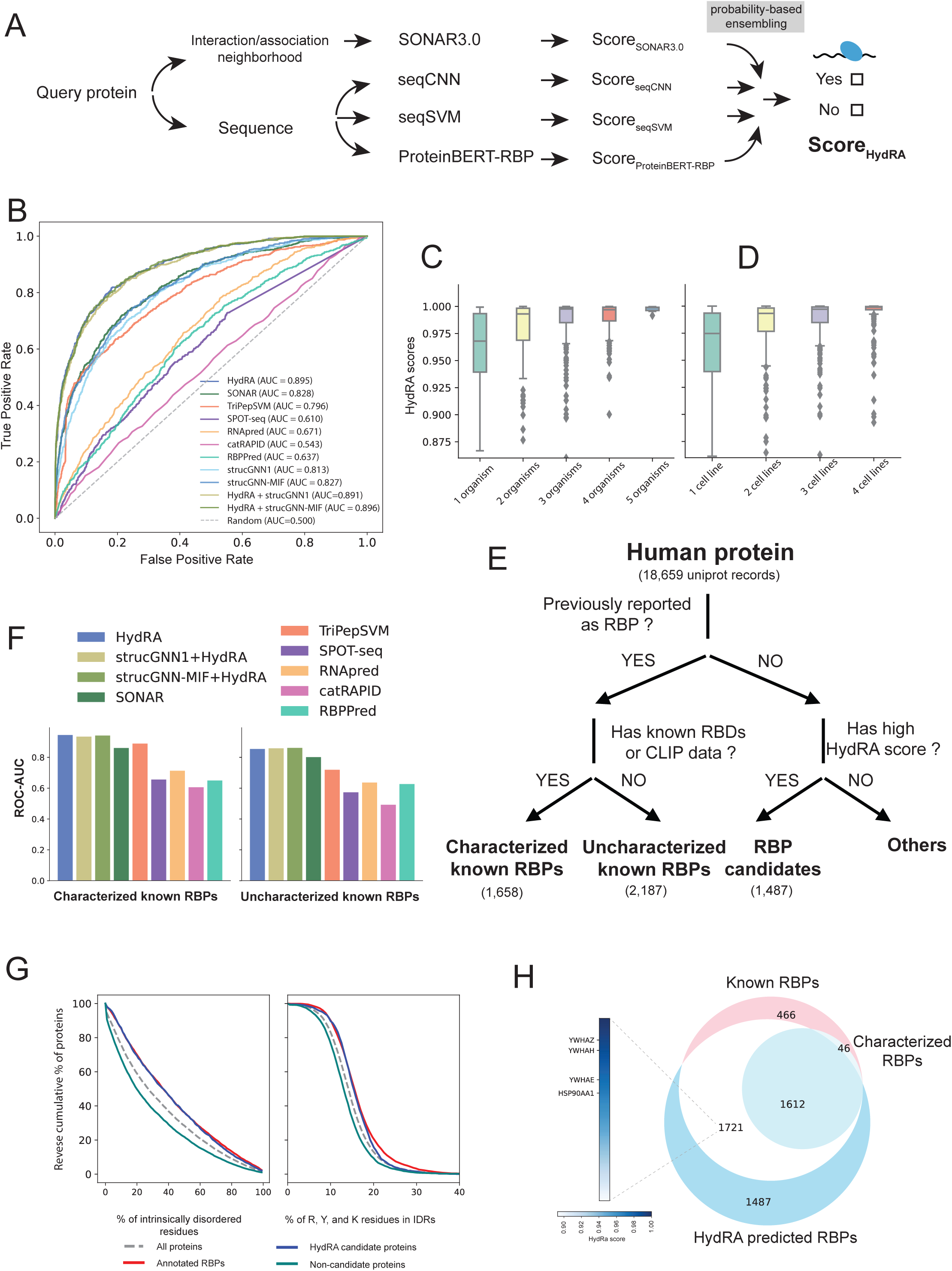
A novel sequence-based approach to recognize RBPs. **A.** The HydRA architecture. The sequence and protein interaction/association neighborhoods of the query protein are fed into four sub-classifiers (i.e. SONAR3.0, seqCNN, seqSVM and ProteinBERT) whose output scores are further integrated by the probability-based ensembling method into final RBP prediction score (Score_HydRA_). **B.** Comparison of current available RBP classifiers according to ROC-AUC analysis within the hold-out test set. Area under curve (AUC) values are indicated for each classifier. **C.** Box plots showing the correlation between the HydRA scores of RBPs and the number of species in which these RBPs have been reported to bind RNA according to the review by Hentze et al.(Hentze et al., 2018b). In the box plots, the lower and upper hinges denote the first and third quartiles, with the whiskers extending to the largest value less than 1.5× the interquartile range. Flier points are those past the ends of whiskers, considered as outliers. **D.** Box plots showing the correlation between the HydRA scores of RBPs and the number of human cell lines in which these RBPs have been reported to bind RNA according to the review by Hentze et al.(Hentze et al., 2018b). **E.** We grouped human proteins into 4 categories. For proteins that are previously reported as known RBPs, we further grouped them to “Characterized known RBPs” and “other known RBPs” based on whether they have characterized RNA-binding domains or CLIP data showing their binding preference. For the rest of proteins, we marked them as “HydRA RBP candidates” if the proteins have HydRA score higher than the score threshold (with false positive rate equal to 0.1), otherwise as “Others”. **F.** We explored the performance of each available RBP classifiers by looking at their recognition capacity on sub-categories of known RNA-binding proteins via ROC-AUC analysis within the test dataset. **G.** We obtained a list of 1,487 candidate RBPs (previously unknown for RNA-binding) with HydRA by setting the false positive rate as 0.1. We compared the physicochemical properties (i.e. the isoelectric points of the proteins, the proportion of intrinsic disordered regions in the protein and the amino acid composition of the intrinsic disordered regions in the proteins) among known RBPs, candidate RBPs, low scored proteins by HydRA (referred to as non-candidate proteins) and all proteins in our protein set. Density curve was plotted to show the difference of each categories in isoelectric points and reverse cumulative frequency curves are showing the difference of each categories in the relative size of intrinsically disordered regions (IDRs) in each proteins and the composition of several RNA-binding-related amino acids in the IDRs of each proteins. **H.** Venn diagram presents the relationship among previously reported RBPs (“Known RBPs”), HydRA predicted RBPs and characterized RBPs. Highly scored uncharacterized RBPs, HSP90AA1, YWHAH, YWHAE and YWHAZ together with YWHAG, are selected for further exploration in this study.

Next, we again grouped known RBPs into characterized and uncharacterized sets (**Figure 3E**) and showed that HydRA had higher recall, specificity, and precision than other classifiers (including HydRA sub-classifiers) for both characterized RBPs and uncharacterized RBPs that lack RNA-binding or -processing related domains and defined RNA targets (ROC-AUC=0.945 and PR-AUC=0.784 for uncharacterized RBPs, ROC-AUC=0.854 and PR-AUC=0.493 for uncharacterized RBPs, **Figures 3F, S3E, S3F**, and **S3G**). Notably, as shown above, TriPepSVM performed well for characterized RBPs but showed more degeneration for uncharacterized RBPs (**Figure 3F**). These results demonstrate that HydRA achieves state-of-the-art predictive power for RBPs either with or without annotated RNA-binding domains, indicating that utilizing protein interaction and association information increases predictive power for RBPs with unknown RNA-binding signatures. This is also evident when we compare the performance of SONAR3.0 with the combination of HydRA’s sub-classifiers for uncharacterized RNA-binding proteins (**Figures S3E** and **S3F**). StrucGNNs displayed slightly better performance than HydRA-seq for recognizing uncharacterized RBPs but showed inferior predictive power for characterized RBPs (**Figures S2D** and **S2E**). Similarly, we observed that incorporating predicted protein structure features into HydRA (referred to as “strucGNN1+HydRA” and “strucGNN-MIF+HydRA”) leads to a slight improvement for recognizing uncharacterized RBPs over HydRA but a slight degeneration on recognizing characterized RBPs (**Figures 3F** and **S3G**). Because these “strucGNNs+HydRA” models do not significantly improve the prediction performance on the overall RBP set (ROC-AUC=0.891 – 0.896, PR-AUC=0.731 – 0.742) shown in **Figure 3B**, we chose not to include strucGNNs in the final HydRA model.

Incidentally, while the HydRA model was only trained with human proteome and human RBP annotations, we observe that this model can also recognize RBPs from mouse, yeast and fly with substantial accuracy producing ROC-AUC values of ∼0.8 and a PR-AUC range from 0.433 – 0.639 (**Figures S4A** and **S4B**). This result indicates the model succeeds in capturing the general RNA-binding associated features across species even just with learning from human RBPs.

We next examined the association between HydRA classification scores and RBP conservation. RBP annotations across multiple organisms (i.e. Homo sapiens, Mus musculus, Saccharomyces cerevisiae, Drosophila melanogaster and Caenorhabditis elegans) and human cell lines (i.e. HeLa, HuH7, HEK293 and K562) collected from a previous study ^29^ revealed that HydRA classifier scores positively correlate with the number of organisms in which a given RBP has been defined (**Figure 3C**). HydRA classification scores also positively correlated with the number of cell-types with validated RNA-binding activity for a given RBP (**Figure 3D**). These results indicate that HydRA scores are clear indicators of the reliability of RBP annotations and establish HydRA analysis as a complementary technique for experimental high-throughput RBP discovery approaches.

HydRA predicts 1,487 human RBP candidates out of the 18,659 proteins that exist in the Mentha-BioPlex network, at a false positive rate of 10% (**Figure 3E, Table S2)**. We observed that HydRA-predicted candidate RBPs are, like known RBPs, significantly enriched in intrinsically disordered regions (IDRs) compared to a background of all human proteins (p < 10^-23^, Mann–Whitney U test). We defined negatively scored predictions as proteins with HydRA scores lower than 50% of the human proteins and these were not statistically different from the background (**Figure 3G**). We also found candidate RBPs had significantly larger proportion of the amino acids arginine, tyrosine and lysine inside intrinsically disordered regions (IDRs), which are biophysical hallmarks of RNA-binding ^29,30^, compared to background (p < 1e-20, Mann–Whitney U test)) and negative predictions (p < 1e-55, Mann–Whitney U test) (**Figure 3G**). These observations demonstrate that HydRA candidate RBPs share similar properties with known RBPs and that mechanisms of IDR-based RNA binding may be widespread among known and unconventional RBPs. Gene ontology analysis of HydRA RBP candidates shows an enrichment for functional categories such as histone modification, DNA replication, and actin cytoskeleton organization, potentially implying RNA-binding functionality for co-transcriptional processes and RNA localization (**Figure S3H**). Uncharacterized RBPs and candidate RBPs can be prioritized by implied biological functions and HydRA predictive scores to explore new RBP classes and RNA-binding landscapes.

### HSP90 is an RBP that binds 3’UTRs of mRNA encoding other heat-shock proteins

HydRA predicts functional RNA-binding for 1,721 uncharacterized RBPs without known RNA-binding domains, RNA-binding functionality, or RNA-targets (**Figure 3H**). To illustrate the utility of HydRA to discover novel RBPs with no previous experimental evidence to support a role in RNA binding, we selected the Heat Shock Protein 90 Alpha Family Class A Member 1 (HSP90AA1) as a high-scoring (HydRA score: 0.968), uncharacterized RBP of substantial biological interest with a classical role as a molecular chaperone that controls cellular homeostasis (**Figure 3H**). While an HSP90 homolog in tobacco plants was found to interact with the genomic RNA of the Bamboo mosaic virus ^31^, there has been no transcriptome-wide characterization of HSP90AA1 as a functional RBP in mammalian systems. We performed eCLIP analysis using V5-tagged HSP90AA1 transiently expressed in HEK293 cells (**Figure S5A**). We obtained highly reproducible binding profiles when compared to control eCLIP of V5 tag expressed without HSP90AA1(**Figure S5B**). Combined HSP90AA1 eCLIP replicate libraries yielded 30,406,215 non-redundant sequenced reads that uniquely mapped to 17,340 annotated protein-coding pre-mRNAs (23,551,600 reads, 77.46%) and noncoding genes such as lincRNAs (929,961 reads, 3.06%), miRNAs (19,498 reads, 0.06%) and tRNAs (9,590 reads, 0.03%). We identified 480 reproducible binding sites (eCLIP peaks) on 225 target genes. Enriched binding motifs were determined from these binding sites with HOMER ^32^ (**Figure S5E**). We found that HSP90 RNA binding sites were highly enriched in the 5’ and 3’ UTRs of target RNAs (**Figures S5C** and **S5D**). Gene ontology analysis of the top HSP90AA1 RNA targets revealed an enrichment for genes related to heat response, RNA metabolism, and G2/M phase cell cycle regulation (**Figure 4A**). Interestingly, most mRNA-targets related to these terms contained HSP90AA1 binding sites within 3’UTR regions. Many of the mRNA-targets related to heat response terms, such as transcripts encoding the HSP70 components HSPA1B and HSPA6, are involved in the HSP90 protein folding pathway (**Figure 4B**). HSP40, HSP70 (composed of subunits that include HSPA1B, HSPA6, and HSPA13), and PTGES3/p23, which itself is more recently discovered as an RBP ^33^, are HSP90 chaperones that help recognize unfolded client proteins, deliver the unfolded proteins to HSP90, and facilitate release of the folded client protein (**Figure 4C**) ^34^. As an orthogonal validation of these HSP90AA1 mRNA-targets, we performed RNA immunoprecipitation followed by qPCR (**Figures S5F** and **S5G**) and observed significant enrichment of mRNAs that encode proteins involved in HSP90 protein folding (HSPA1B, HSP6, HSP40) and RNA metabolism (MBNL1 and CNOT1) but not the mRNAs encoding ACTIN or GAPDH. These results reveal a highly specific RNA-binding landscape for HSP90AA1 that likely regulates chaperone responses to cellular stress.

**Figure 4.**
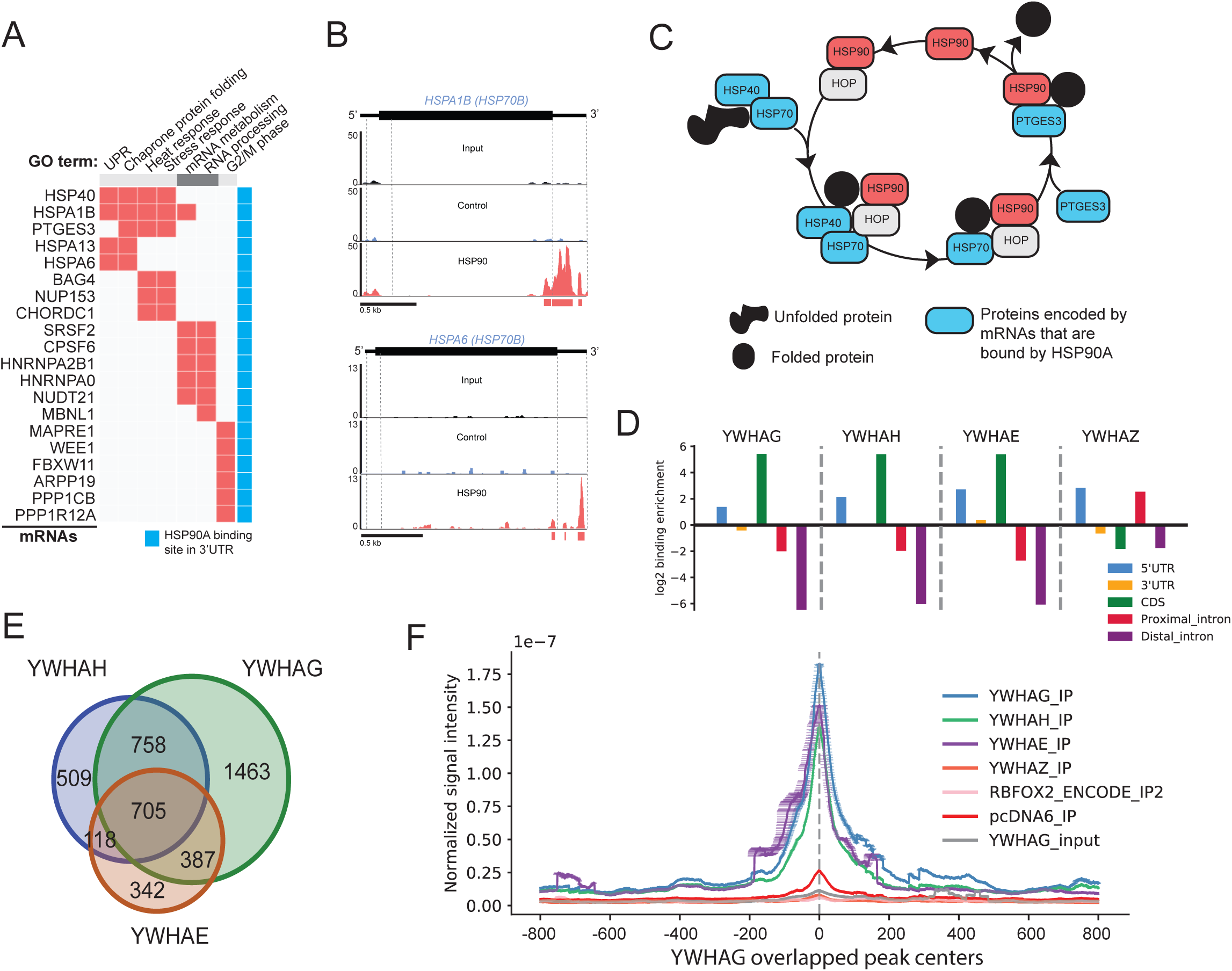
eCLIP analysis discovers HSP90 as an RBP that binds mRNA encoding other heat-shock proteins, and YWHA (14-3-3) family that binds RNAs with similar binding preference across the transcriptome. **A.** Gene ontology enrichment analysis for the 100 top HSP90 RNA targets using the Enrichr Gene Ontology enrichment tool identified GO terms related to heat response, mRNA metabolism/processing, and cell cycle G2/M phase. RNAs related to the GOs are indicated by the orange box. The blue box indicates the RNAs that are bound by HSP90A in their 3’ UTR. **B.** Genome browser shots of HSPA1B (top) and HSPA6 (bottom), which both belong to the HSP70B complex, showing an enrichment of sequening reads in the 3’ UTRs of both mRNAs in the HSP90A eCLIP-seq library but not in the input library or Control eCLIP-seq library. **C.** Shematic of the HSP90 protein folding pathway shows that many of the constituients in this pathway are RNA targets of HSP90A. **D.** Binding preference of YWHAG, YWHAH, YWHAE, and YWHAZ on different genomic regions measured by log2-transformed region enrichment E_region_ (see Methods). The following genomic regions of a transcript are shown: 5’UTR (blue), 3’UTR (yellow), coding sequence (CDS, green), proximal introns (red), and distal intron (purple). **E.** The venn diagram for the overlap of RNA targets of YWHAG, YWHAH and YWHAE. **F.** Read density plot around the binding peak center of YWHAG across the transcriptome. The read density of the eCLIP IP samples of YWHAG, YWHAH, YWHAZ, RBFOX2, vector control and the size-match input samples of YWHAG is shown.

### Whole-transcriptome characterization of YWHAG/H/E/Z as RNA-binding proteins

YWHA (14-3-3) is a family of multifunctional proteins that work as phosphor-binding adapters, scaffolding proteins, and protein chaperones. They are known for their involvement in the critical processes of cancers (such as apoptosis, cell cycle progression and autophagy) and neurological disorders ^35–39^. Interestingly, all the members of this family were assigned with high HydRA scores (**Figure 3H**); and while they have been identified as RBP candidates in one or more high-throughput screens ^5,6,30,40,41^, genome-wide RNA targets and RNA-binding functionality have not been confirmed for this protein family.

Here, we present a transcriptome-wide characterization of RNA-binding sites of YWHAH, YWHAG, YWHAE and YWHAZ proteins with eCLIP technology using V5-tagged proteins transiently expressed in HEK293 cells (**Figure S6B**). With the peak calling algorithm CLIPper, we identified 11,187 binding sites (eCLIP peaks) over the transcriptome with stringent threshold (see Methods) shared by YWHAG eCLIP replicates, 4,506 common binding sites in YWHAH replicates, 2,939 common binding sites for YWHAE and 308 common binding sites for YWHAZ (**Figure S6A**) with their enriched binding motifs on the RNA targets determined by HOMER ^32^ (**Figure S6E**). Three of the four proteins (viz. YWHAG, YWHAH and YWHAE) have their binding sites enriched in the protein coding (CDS) and 5’UTR regions of mRNA molecules and under-presenting in intronic regions, while the binding sites of YWHAZ are slightly enriched in 5’UTR and proximal intronic (defined as the intronic regions that are with the distance < 500 bp from exon-intron junction) regions (**Figures 4D** and **S6C**). Furthermore, we found high overlap in the RNA targets of YWHAG, YWHAH and YWHAE, where about 70% of the RNA targets of YWHAH and YWHAE are shared with YWHAG (**Figure 4E**). The locations of the binding sites of YWHAH and YWHAE are also highly correlated with YWHAG as we plotted the density of sequence reads around YWHAG binding sites (**Figure 4F**) while YWHAZ, RBFOX2 and negative controls (i.e. the reads from size-match INPUT sample of YWHAG and empty vector) shows no significant binding behaviors around YWHAG’s binding sites. An example of YTDC1, a regulator of alternative splicing, was shown with genome browser tracks (**Figure S6D**). GO analysis of the common RNA targets of these three proteins shows an enrichment in RNA splicing and translational regulation process (**Figure S6F**). All of these results suggest that YWHAG, YWHAH and YWHAE are probably working synergistically or antagonistically on RNAs and may participate in the regulation of RNA splicing and translation processes.

### Occlusion Mapping interprets sequence-based RBP prediction and highlights functional elements in RBPs

In computer vision, occlusion analysis is widely used to interpret the validity of machine learning models. This analysis tests the impact of occluding a sub-region of the image on the inference step to determine if this sub-region is essential for the model to recognize the identity of the image. We carried out an equivalent procedure for RBP classification with the sequence-based component of HydRA, referred to here as Occlusion Mapping (**Figure S7A**). Occlusion Mapping eliminates a window of twenty consecutive amino acids from the protein sequence in the inference step and compares the model output score with the score from the intact protein sequence i.e. the difference of the scores is denoted by Δscore after accounting for protein lengths and model-specific score distributions to determine whether or not the sequence is pertinent to the protein’s ability to bind RNAs (**Figure S7A**, details in Methods). A negative Δscore indicates that the corresponding region of the occluded window contributed positively to the RNA binding activity. We determined a region’s (occlusion window) predicted ability to bind RNA from the p-values generated from the Δscores of a that specific region. We define continuous stretches of occlusion windows with p-values < 0.05 as “predicted RNA-binding regions” (pRBRs) and with p-values < 0.001 as stringent pRBRs.

We display the occlusion maps for eight well known RBPs that are involved with distinct roles in RNA processing ranging from splicing to mRNA decay and translation (RBFOX2, ZFP36, ATXN2 and RPLP0 in **Figure 5A**; and PUF60, LSM11, EIF4E and YBX2 in **Figure S8A**). In RBFOX2 (**Figure 5A**), we observed a contiguous area of negative Δscores with stringent pRBRs that correspond well with its known RNA recognition motif (RRM). We also found a contiguous area with weaker but significant occlusion signals (pRBRs) (p<0.05**, Figure 5A**) in a Calcitonin gene-related peptide regulator C terminal domain (Fox-1_C), which has been described to regulate alternative splicing events by binding to 5’-UGCAUGU-3’ elements ^42^. Similarly, we observed two peaks of stringent pRBRs in the two zinc finger domains (zf-CCCH) described to bind RNA in ZFP36 ^43^. Since this zinc-finger domain consists of 22 amino acids, this demonstrates that our sequence-based classifier with occlusion map can recognize RNA-binding protein regions at high resolution. There are also strong correspondences between Δscore peaks within pRBRs/stringent pRBPs and diverse annotated RNA-binding domains found in ATXN2, RPLP0, PUF60, LSM11, EIF4E and YBX2 (**Figures 5A** and **S8A**).

**Figure 5.**
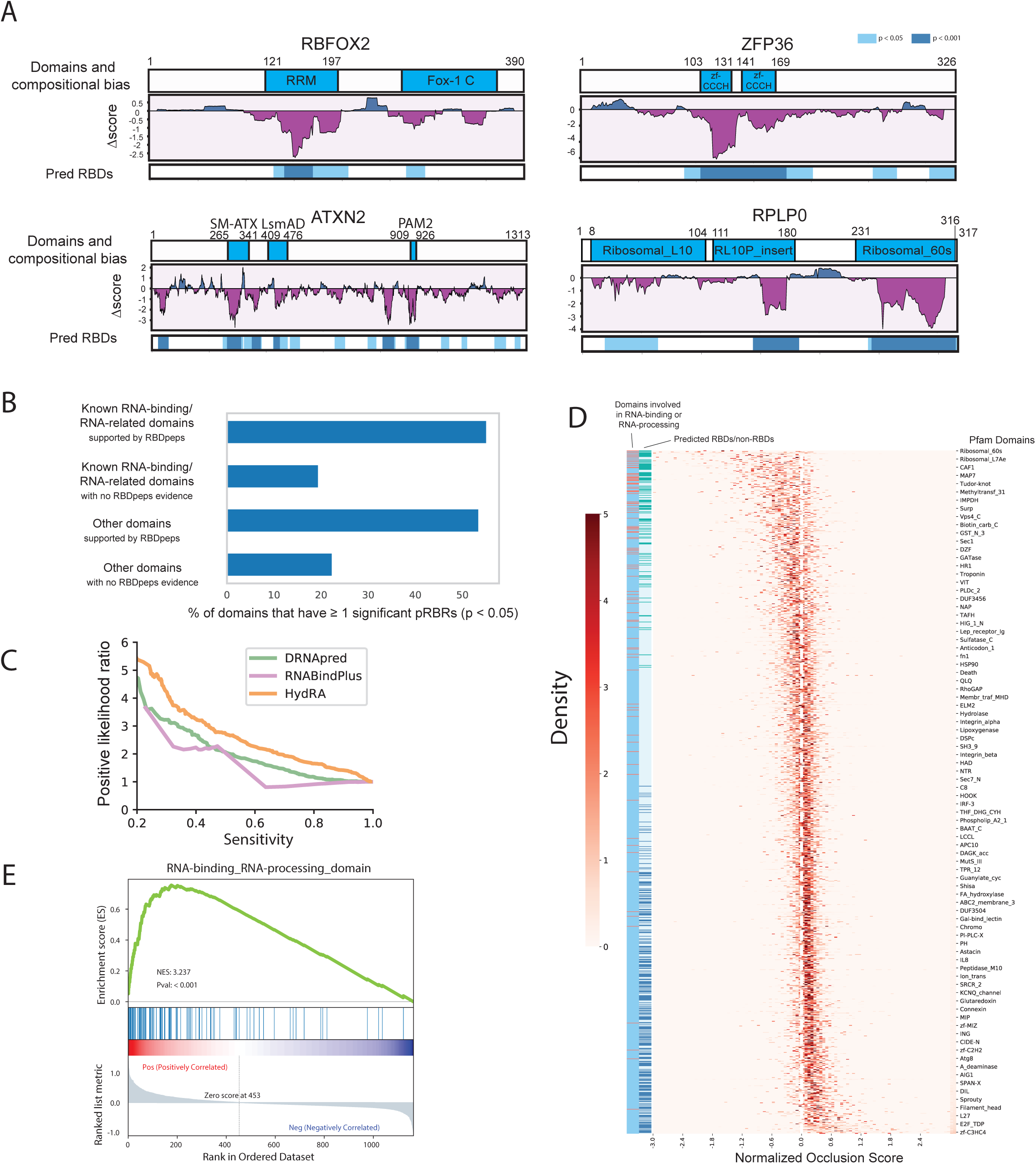
Occlusion map enables model interpretation of HydRA Sequence Classifier and novel RNA-binding domain discovery. **A.** Occlusion Maps of several known RBPs (including RBFOX2, ZFP36, ATXN2 and RPLP0). The coordinates of protein domains, compositional bias and RBDpeps are shown. Protein regions that have Δscore < 0 are colored with purple, while those with Δscore > 0 are colored with blue in the middle panel. The regions with significant p values are also marked in the bottom track: skyblue for p<0.05 and dark blue for p<0.001. **B.** We grouped the protein domains in known RBPs to four categories: RNA-binding domains (RBDs) that are supported by RNA-binding peptide studies, RBDs that lack support from current RNA-binding peptide studies, other domains with and without evidence from RNA-binding peptide studies. The bar plot represents the percentage of the domains within each category that have ≥ 1 occluders with significant negative Δscores (p-value < 0.05). **C.** To evaluate the domain-level RBD detection performance, the positive likelihood ratio at different sensitivity level were visualized for baseline algorithms for RNA binding-residue prediction (DNAPred and RNABindPlus) and HydRA occlusion map. **D.** A heatmap showing the distribution of occlusion scores for each human protein domains (with more than 5 human protein members). Domains are sorted by their average occlusion scores from small to large. Protein domains that are already known for RNA-binding or involvement in RNA processing are marked in red in the first annotation bar (on the right side) of the heatmap. The second annotation bar shows the predicted RBD (mean > 0 and FDR adjusted p-value < 0.05, one-sample t-test) in green and predicted non-RBD (mean > 0 and FDR adjusted p-value < 0.05) in dark blue. **E.** A GSEA plot showing that the set of RNA binding or RNA-processing related domains are enriched in the top part of the ranked list of human domains shown in B. The top portion of the plot shows the running Enrichment Score (ES) for the domain set of RBD/RNA-processing as the analysis walks down the ranked list. The middle portion of the plot shows where the domains with RNA--binding or RNA processing function appear in the ranked list of human domains. The bottom portion of the plot shows the value of the ranking metric as you move down the list of ranked genes.

In general, we discover that annotated RNA-binding or -processing related domains are 2.5X more likely to contain pRBRs (p<0.05) than non-RNA-binding regions of the human proteins (**Figure 5B**) as evaluated by positive likelihood ratios (see Methods), while the ratio increases to 4.5X when observing stringent pRBRs (p<0.001) (**Figure S8D**). Additionally, the high-confidence RBDs, which are supported by experimentally identified RNA-binding peptides (referred to as RBDpeps) studies ^6,44–46^, are more enriched by pRBRs and stringent RBRs than the RBDs or RNA processing domains without no experimental evidence (**Figure 5B and S8D**), indicating the robustness of Occlusion Map to detect bona fide RNA binding domains. In addition, although HydRA was not trained explicitly with RNA-binding annotations, it achieves slightly better performance on the domain-level characterization of RNA-binding elements than the baseline tools in the field, i.e. RNABindPlus and DRNApred, which were originally designed to predict RNA-binding residues in RBPs (**Figure 5C**). To discover potential new RNA-binding domains (RBDs) from the perspective of HydRA, we obtained occlusion score distributions for each human protein domain from the human protein sequences and visualized them as heatmaps (**Figures 5D, S8B**, and **S8C**), where domains are sorted by the mean of their occlusion score distributions. One-sample t-test and Benjamini-Hochberg method for multiple test correction recognized the domains that have Δscores significantly less than 0 (predicted RBDs) or larger than 0 (predicted non-RBDs). Domains with greater than 5 human protein members are included, 1,165 domains in total. We observed the majority of the RNA-binding and RNA-processing related domains (including LSM, CSD, RRM, KH, zf-CCCH, etc.) appear in the top 100 of the sorted domain list with significant p-values (**Figure S8B**). A large majority of the bottom 100 domains are unrelated to RNA-binding or RNA-processing except RNA polymerase Rpb2 domain 7, which is known for DNA-binding^47^ and is not statistically significant (FDR-adjusted p-value = 0.102) (**Figure S8C**). GSEA analysis also shows most of the known RBDs and RNA processing related domains are enriched in the top panel of this sorted list (normalized enrichment score = 3.237, p value < 0.001, **Figure 5E**). Furthermore, many of the top binding domains with significant p-values (76 out of 104 domains, FDR-adjusted p < 0.05) are not canonical RNA-binding domains or known to be related to RNA processing, thus we obtained a list of 76 new RNA binding domains from this analysis (**Table S3**).

### Validation of RNA Binding Activity and RNA-binding elements of Candidate RBPs

We next evaluated HydRA’s ability to predict RNA binding activity at the region level for a selection of candidate RBPs harboring domains with significant Occlusion delta scores. We selected candidates with representative domains, such as zinc-finger (zf) and EF hand domains, that received top RBD predictive scores from occlusion map analysis (**Figure S8B**). Individual occlusion mapping across the 5 candidate RBPs INO80B, PIAS4, NR5A1, ACTN3, and MCCC1 revealed specific domain regions that are predicted to bind RNA (**Figure 6A**). We performed eCLIP on these V5 tagged RBP candidates, with and without these occlusion-predicted RNA-binding elements and assessed transcriptome-wide RNA targets (**Figure 6B**). In total, we obtained 48 eCLIP libraries (including SMInput) for these 5 candidate RBPs. Each library was sequenced to 5.7 – 57.0 million reads, of which ∼ 0.42 – 16.0 million reads mapped uniquely to the human genome. We used a stringent peak-calling workflow that included both size-matched and V5-tag-only inputs as well as both Clipper^48^ and ChIP-R reproducible peak-calling algorithms^49^ (**Figure S9C**), which identified between 110 (INO80B) and 2,849 (ACTN3) RNA-binding peaks per candidate (**Figure 6D**). Ontology analysis of bound targets revealed that INO80B bound genes are involved in translation and have binding sites enriched in 5’UTR regions, PIAS4 bound genes are involved in signaling and differentiation and have sites in proximal intronic regions, NR5A1 bound genes are involved in mitosis with CDS binding sites, ACTN3 bound genes are involved in ER-stress with sites in 5’UTR regions, and MCCC1 bound genes are involved in cell morphology and polarity with sites broadly across 5’UTR, CDS and 3’UTR regions (**Figures 6C** and **S9D**). These RNA-binding profiles suggest roles for these candidates in regulating the cytoplasmic localization and/or translation of target genes in the cases of UTR and CDS binding, and alternative splicing regulation in the case of intronic binding.

**Figure 6.**
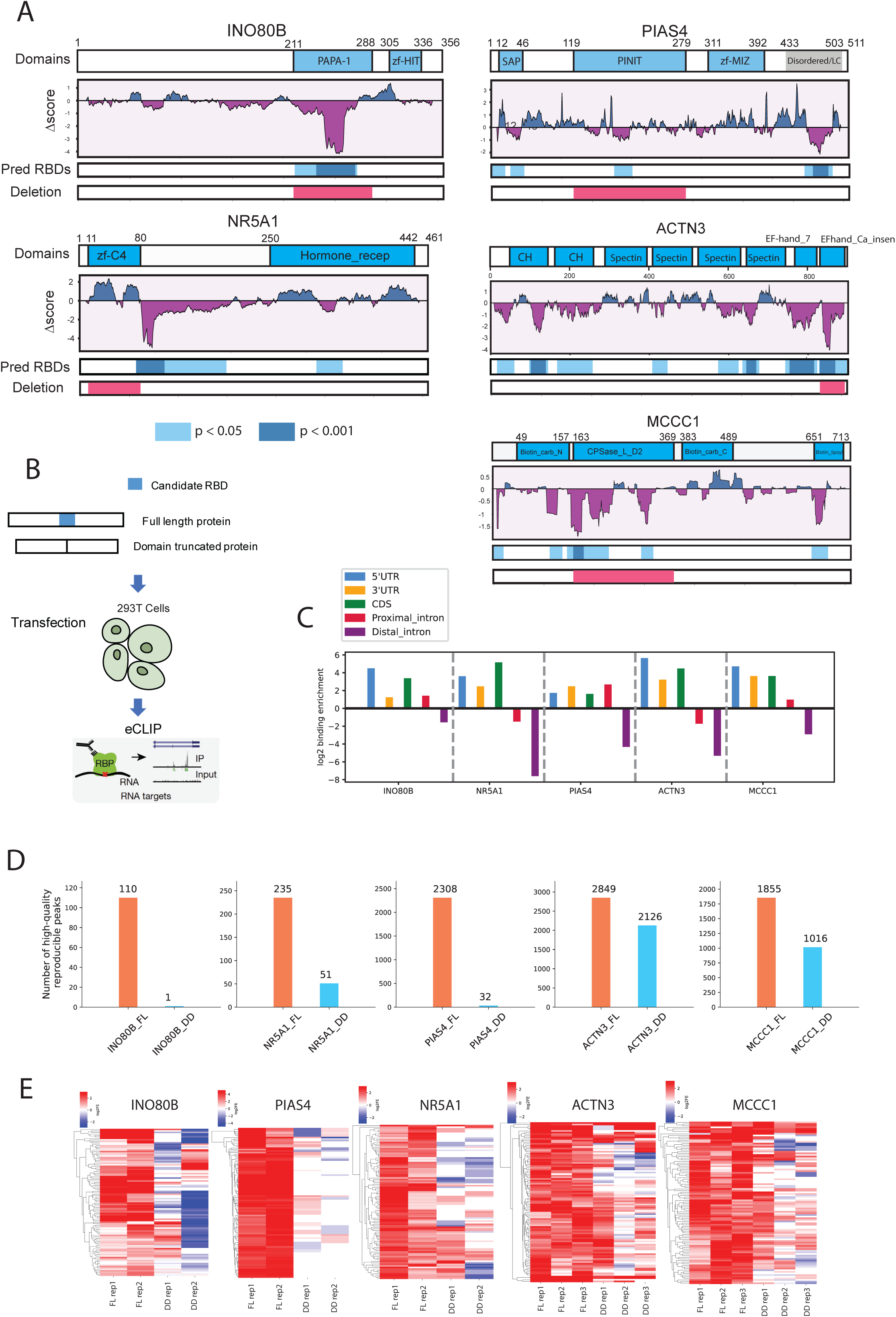
Novel RBPs and functional regions related to RNA-binding were predicted using HydRA and experimentally validated. **A.** Occlusion Maps of INO80B, NR5A1, PIAS4, ACTN3, and MCCC1. The coordinates of protein domains and compositional bias are shown. Protein regions that have Δscore < 0 are colored with purple, while those with Δscore > 0 are colored with blue in the middle panel. The regions with significant p values are also marked in the bottom track: skyblue for p<0.05 and dark blue for p<0.001. **B.** The workflow to experimentally validate candidate RBPs and its corresponding predicted functional domain for RNA binding behavior. Original and truncated protein (with predicted functional domain deleted) were expressed in HEK293T cells followed by applying eCLIP experiments to explore their RNA-binding behaviors. **C.** Binding preference of INO80B, NR5A1, PIAS4, ACTN3, and MCCC1 on different genomic regions measured by log2-transformed region enrichment E_region_ (see Methods). The following genomic regions of a transcript are shown: 5’UTR (blue), 3’UTR (yellow), coding sequence (CDS, green), proximal introns (red), and distal intron (purple). **D.** The bar plots showing the number of reproducible eCLIP peaks for full-length and truncated protein of INO80B, NR5A1, PIAS4, ACTN3, and MCCC1. **E.** The heatmap showing the binding strength (measured by log transformed fold-enrichment) across full-length (FL) and truncated (DD) protein samples on the 100 most significant peaks found in the full-length proteins.

Removal of putative RNA-binding domains with significant delta-occlusion scores for these candidates did not influence expression of V5-fusions as assessed by western blot analysis (**Figure S9A**). However, absence of these predicted domains did result in substantially higher PCR cycle requirements (eCT values) to obtain eCLIP libraries from immunoprecipitated RNA (**Figure S9B**), indicating the importance of the removed domains for RNA-binding activity. By comparing the eCLIP peaks of the full-length proteins (FL) and domain-deleted proteins (DD) (**Figure 6D**), we observed that these putative RNA-binding domain deletions coincided with a global loss of 2,779 eCLIP peaks on 2,089 targets (out of 2,849 peaks on 2,123 targets) for ACTN3, and 1,820 peaks on 1,576 targets (out of 1,855 peaks on 1,597 targets) for MCCC1, 110 peaks on 151 targets (out of 110 peaks on 151 targets) for INO80B, 2,308 peaks for eCLIP peaks on 1,187 targets (out of 2,308 peaks on 1,187 targets) for PIAS4, 232 peaks on 235 targets (out of 235 peaks on 238 targets) for NR5A1. Analysis of the top 100 binding peaks for these RBP candidates upon candidate domain deletion revealed highly reproducible reduction in RNA-binding across these high confidence binding sites (**Figure 6E**). This drastic loss of RNA-binding upon domain deletion demonstrates the high-resolution accuracy with which HydRA predicts RNA-protein interactions. Together these results validate the power of the HydRA workflow for both RNA-binding protein (gene-level) and RNA-binding domain (amino-acid level) discovery.

## DISCUSSION

The rapidly growing importance of RBP biology and function in development and disease necessitates a comprehensive RBP census. Recently, several high-throughput biochemical approaches have expanded the number of putative RBPs in several cell-lines. In parallel, several computational approaches (RBP classifiers) have been developed to complement these RNA-interactome assays and achieve higher-accuracy RBP discovery. The resultant expanding RBP catalog presents a growing diversity of “unconventional” RBPs that do not contain well-defined RNA-binding domains^29^. These unconventional RBPs present obstacles to current RBP classifiers (see **Figures S2A, S2B, 3B** and **S3C**) resulting in impaired accuracy. To overcome these obstacles, we developed HydRA as a learning based RBP classifier, that recognizes RNA-binding proteins based on both external protein context (using protein-protein interaction networks) and internal amino acid sequence signatures. HydRA employs this external-context:internal-pattern combination analysis to achieve unprecedented sensitivity, specificity and precision for predicting protein-RNA interactions. The use of deep learning enabled our Occlusion Map application that interprets the HydRA model to predict RNA-binding amino acid sequence regions within proteins without additional training or prior RNA-binding domain information. A proteome-wide HydRA search produced extensive candidate lists of both RNA-binding proteins and RNA-binding domains. We experimentally characterized the transcriptome-wide RNA-binding sites of five new RBPs identified by both HydRA and RNA-interactome capture studies, adding new insights into the functions of these proteins. We further experimentally validated the RNA-binding capability of five HydRA-predicted RBP-candidates at amino acid level resolution, thereby elucidating previously unknown, conserved RNA-binding domains.

In previous work, we proposed a PPI-based RBP classification algorithm, SONAR, which optimized performance across the proteomes of several species ^12^. One drawback of the SONAR approach is the emergence of false-negative predictions when the selected PPI information is limited (**Figures 1A** and **1B**). HydRA alleviates this limitation with an upgraded protein-context component (SONAR3.0) which inherits the main features of SONAR but markedly improves performance by (1) employing a more comprehensive collection of experimental PPI data in place of any single high-throughput PPI study (2) incorporating more context information with a sub-component retrieving features from functional protein association network; (3) penalizing predictions inferred from inadequate PPI or functional association data. We combined SONAR3.0 with an added sequence-based component (HydRA-seq) aimed at recognizing new RBPs by detecting sequence-level patterns overrepresented in known RBPs. In this component, the RBP-exclusive sequence patterns are modelled and processed with a support vector machine model (seqSVM), a convolutional neural network (seqCNN), and an attention-based protein language model (ProteinBERT-RBP) respectively to infer RNA-binding capacity for any given protein. Using a probability-based ensemble approach, we integrated the predictions of these three classifier pieces to obtain an ensemble classifier that achieves higher performance than each individual classifier alone, especially in the recognition of RBPs without annotated RNA-binding domains. The HydRA sequence-based component (HydRA-seq) not only outperforms current state-of-the-art sequence-based RBP classifiers (such as TriPepSVM) but also enables the first domain-level RNA-binding predictions using an Occlusion Map application. Surprisingly, without including RNA-binding information (such as RBD annotations and RNA-binding residues) in the training process, the HydRA-powered Occlusion Map outperforms the baseline RNA-binding residue predictors in recognizing RNA-binding domains. Notably, the SONAR3.0 component of HydRA outperforms the sequence-based component and all other published RBP classifiers, all of which are sequence based. This PPI-component behavior suggests that a large portion of the RBP catalog, particularly unannotated RBP candidates, lack common sequence-level patterns that can be easily captured by current machine learning models, and that protein-protein interaction and functional protein association data provides highly informative patterns for classifying RBPs. Integrating the output from these two HydRA components yields an overall performance improvement compared to individual components, revealing that RNA-binding proteins are most accurately predicted when both external protein context and internal sequence signatures are leveraged.

HydRA scores are highly correlated with the number of experimental studies that identify a given RBP as cross-linkable to RNA. This correlation demonstrates that HydRA can serve as a valuable complement for high-throughput RBP discovery approaches by scoring high probability candidates and reducing the false discovery rates. The sensitivity of the Occlusion Map approach from the sequence-based component of HydRA potentially extends HydRA’s applicability to *in silico* perturbation analyses to evaluate how protein sequence variations influence RNA-binding capacity. Moreover, since the HydRA framework does not depend on prior knowledge of RBPs (such as RNA-binding domains), this framework can be easily transferred and applied to RBP prediction in other species and even other protein function prediction problems with appropriate training dataset.

We evaluated a representative set of HydRA-predicted RBPs by eCLIP to characterize transcriptome-wide targets. The previously uncharacterized RBP HSP90AA1 preferentially binds to transcripts encoding heat response proteins involved in the HSP90 protein folding pathway, and the uncharacterized RBP members of the 14-3-3 (YWHA) family of proteins targets genes involved in RNA splicing and translation processes. The 14-3-3 domain shared by the YWHA family members was predicted by occlusion analysis to contribute to RNA-binding activity, and thus represents a previously uncharacterized RNA binding domain. Enhanced CLIP binding profiles indicated potential synergistic or antagonistic cooperation of YWHAG/E/H binding to RNA targets.

Many new candidate RBPs predicted by HydRA contain significant occlusion delta scores in domains not previously known to confer RNA-binding activity. These domains represent valuable additions to the existing catalogue of characterized RNA-binding domains. To verify RNA-binding for some of these candidate proteins and domains, we performed eCLIP using full-length and putative RNA-binding domain-deleted expression constructs. This approach demonstrated RNA-binding that was dependent on Occlusion Map-predicted domains including the PAPA-1 domain of INO80B, the PINIT domain of PIAS4, the zf-C4 domain NR5A1, the EF-hand-CA-insen domain ACTN3, and the CPSase_L_D2 domain of MCCC1. With future validation efforts, HydRA predicted RBPs from this study (with FPR=0.1) would add 1,487 new RBPs, with potentially a hundred new RNA-binding domains, to the catalog of annotated human RBPs.

In conclusion, the HydRA algorithm is a powerful protein context and sequence-based method for the *de novo* identification of candidate RBPs and RNA binding regions. In silico analysis with HydRA accurately predicts the properties of known and novel RBPs including functional domains. Future efforts to validate and characterize the extensive list of HydRA predicted RBP candidates and novel RNA-binding domains will deepen our understanding of RBP roles in diverse biological processes. HydRA suggests a general machine learning framework for integrating protein-protein interaction and protein sequence information for the predictions of RBPs, and potentially other protein classes, whenever both features contribute substantially to functional activity. Lastly, given the accurate knowledge learned from RBPs, HydRA, particularly its generative component (i.e. ProteinBERT-RBP), also provide an opportunity for novel RBP design, which could be coupled with current protein design workflow and other protein properties classifiers to design novel RBPs that are more suitable for specific biological tasks.

## AUTHOR CONTRIBUTIONS

W.J. and G.W.Y conceived the study. W.J. developed HydRA, wrote the software, tested the software, collected data, analyzed data and visualized results. K.W.B., K.K., G.W.Y., W.J. and J.S.X. designed wet-lab validation experiments. K.W.B., K.K., S.S.P, H.Q.T., M.L.G, M.M and J.A. carried out the experimental work. W.J., G.W.Y and L.W. designed computational validation experiments. W.J. and B.H. carried out the computational validation experiments. W.J., K.W.B. and K.K wrote the original manuscript draft. W.J., K.W.B., K.K., G.W.Y., S.S.P. and K.R. reviewed and edited the manuscript. G.W.Y. acquired the funding and supervised the study.

## ACKNOWLEDGMENTS

G.W.Y. is supported by NIH R01 HG004659, U24 HG009889 and an Allen Distinguished Investigator Award, a Paul G. Allen Frontiers Group advised grant of the Paul G. Allen Foundation.

## SUPPLEMENTAL FIGURE TITLES AND LEGENDS

**Figure S1: Related to Figure 1.**
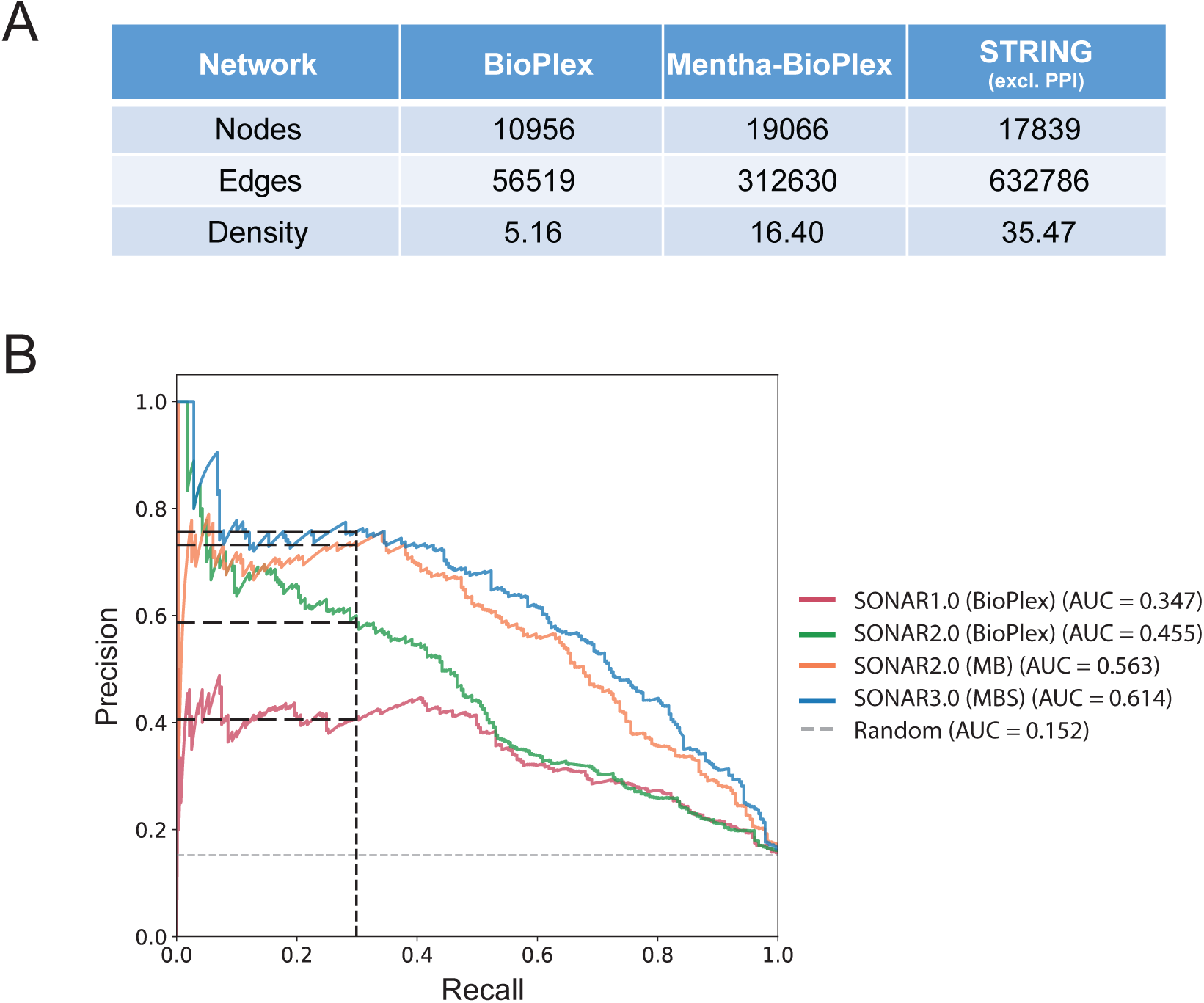
Protein Interaction and Association based (SONAR3.0) classifier presents upgraded RBP classification performance from SONAR. **A.** PPI network summary for BioPlex, Mentha-BioPlex and STRING. Notably, only predicted PPIs of STRING network were used in this study. **B.** Precision-recall AUC analysis of classifiers using SONAR1.0 with BioPlex network, SONAR2.0 with BioPlex network, SONAR2.0 with MB network and SONAR3.0 with MBS networks. The grey dashed line represents the expected performance of random models and the black dashed lines mark the precision of each classifier at a recall of 30%.

**Figure S2: Related to Figure 2.**
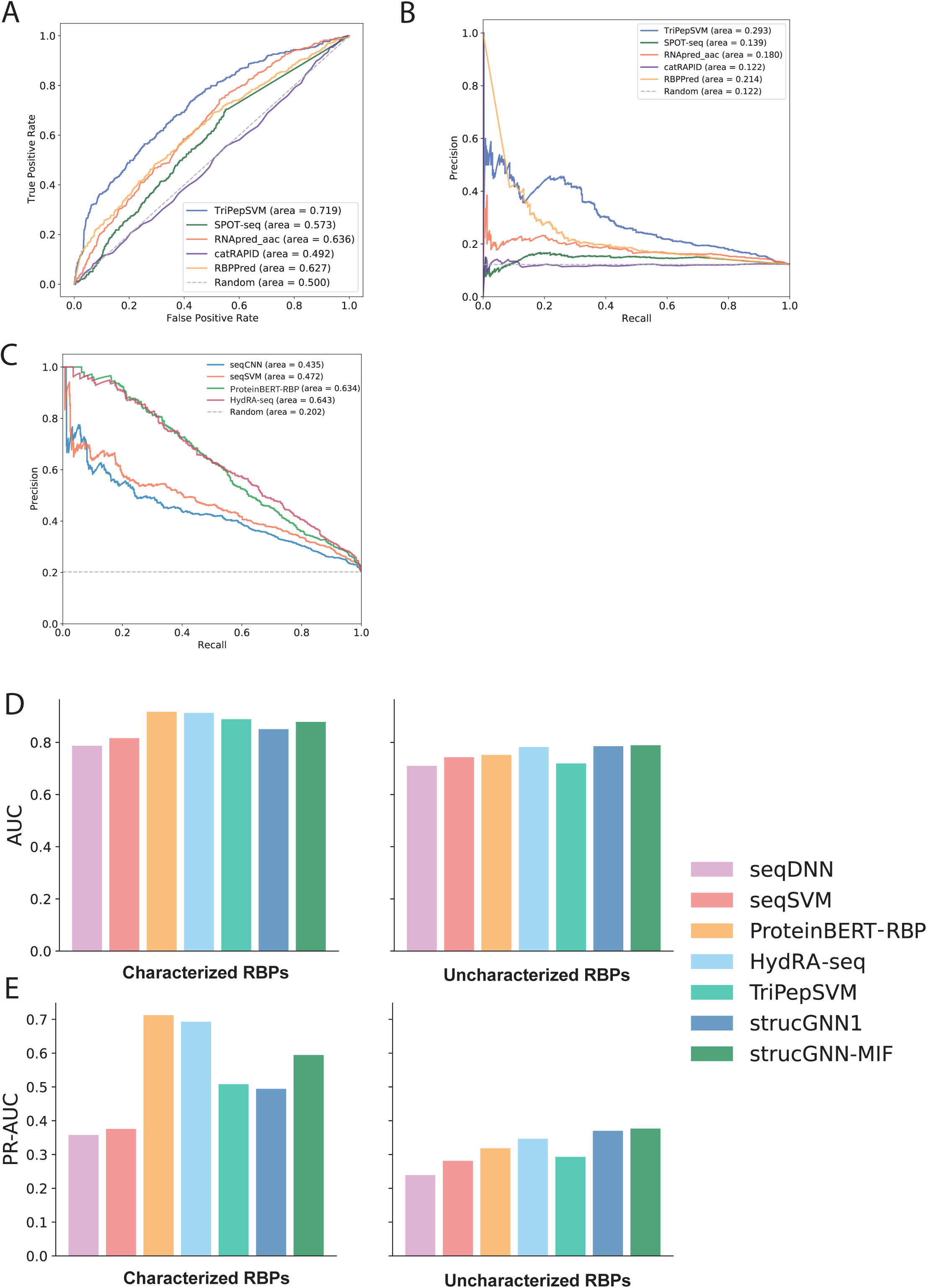
The performance of our sequence-based classifiers. **A.** ROC-AUC analysis showing the performance of TriPepSVM, SPOT-seq, RNApred, catRAPID and RBPPred on recognizing “uncharacterized known RBPs” in our test dataset. **B.** Precision-recall analysis showing the performance of TriPepSVM, SPOT-seq, RNApred, catRAPID and RBPPred on recognizing “uncharacterized known RBPs” in our test dataset. **C.** Plot of the precision-recall analysis of seqCNN, seqSVM, ProteinBERT and Sequence Classifier (Combined). The theoretical random performance is shown as a dashed line. **D-E.** We explored the performance of sequence-based HydRA classifiers by looking at their recognition capacity on sub-categories of known RNA-binding proteins (i.e. “characterized” and “uncharacterized” known RBPs) via ROC-AUC and precision-recall analysis within the test dataset. The performance of TripPepSVM and strucGNNs are also shown in the plots.

**Figure S3: Related to Figure 3.**
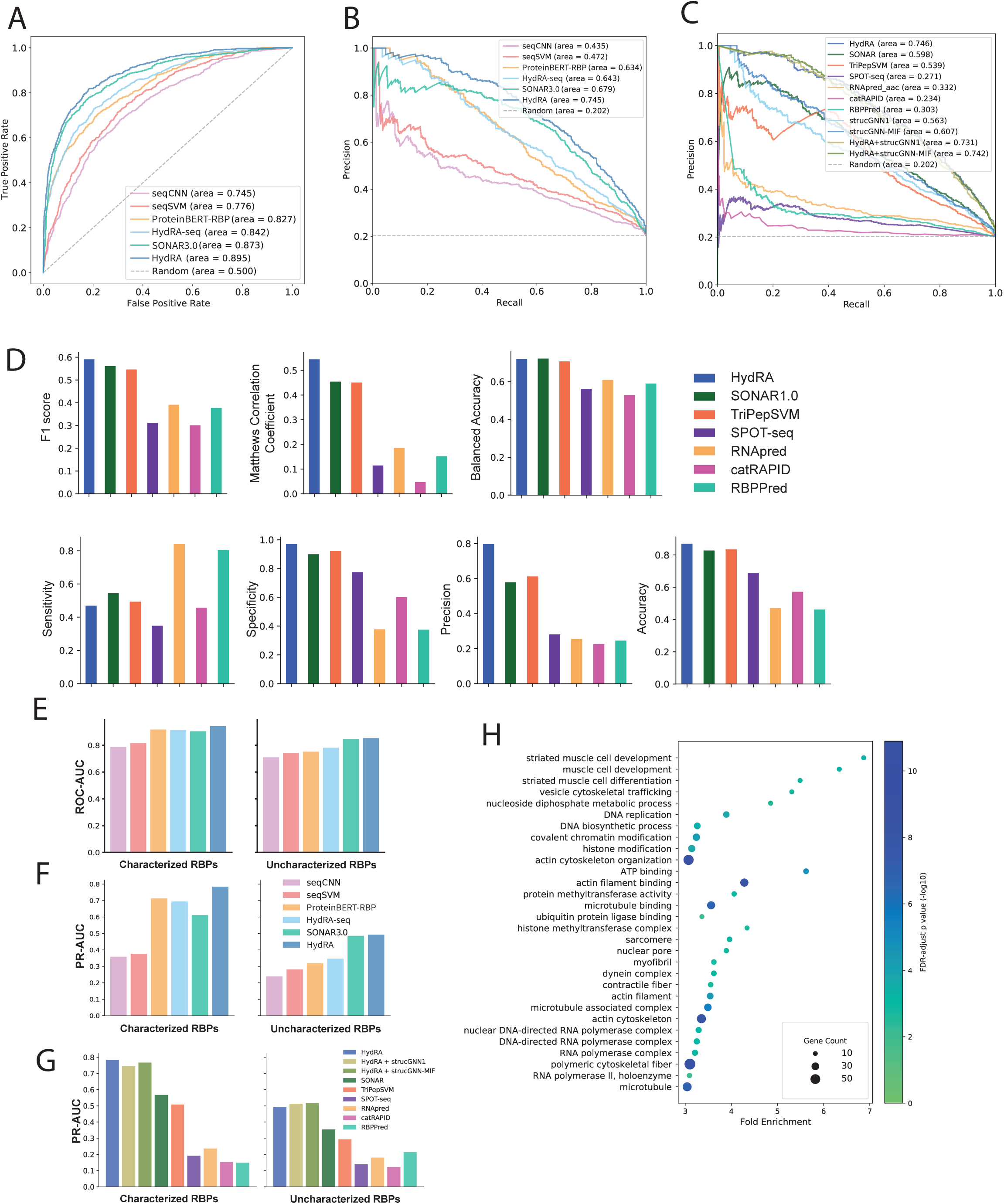
Comparing classification performance of RBP classifiers. **A.** Plot showing the ROC-AUC analysis of each sub-classifier (component) in HydRA. The theoretical random performance is shown as a dashed line. **B.** Plot showing the precision-recall analysis of each sub-classifier (component) in HydRA. The theoretical random performance is shown as a dashed line. **C.** The plot of precision-recall analysis that compares the classification performance of current RBP classifiers (HydRA, SONAR (trained and test with Mentha-BioPlex network), TriPepSVM, SPOT-seq, RNApred (aac mode), catRAPID signature and RBPPred). The theoretical random performance is shown as a dashed line. **D.** Bar plots showing the classification performance of current RBP classifiers with metrics such as sensitivity, specificity, precision, accuracy, Matthews correlation coefficient (MCC), balanced accuracy and f1 score. The classification score threshold for each classifier was defined by their original papers (or the default setting of the software), while the classification score threshold of SONAR and HydRA was determined by keep the false positive rate as 0.1 (as mentioned in the main text). **E-G.** We explored the performance of each sub-classifiers of HydRA, strucGNN+HydRA and other available RBP classifiers by looking at their recognition capacity on sub-categories of known RNA-binding proteins (i.e. “characterized” and “uncharacterized” known RBPs) via ROC-AUC and precision-recall analysis within the test dataset. **H.** The enriched GO terms in HydRA candidate RBPs.

**Figure S4: Related to Figure 3.**
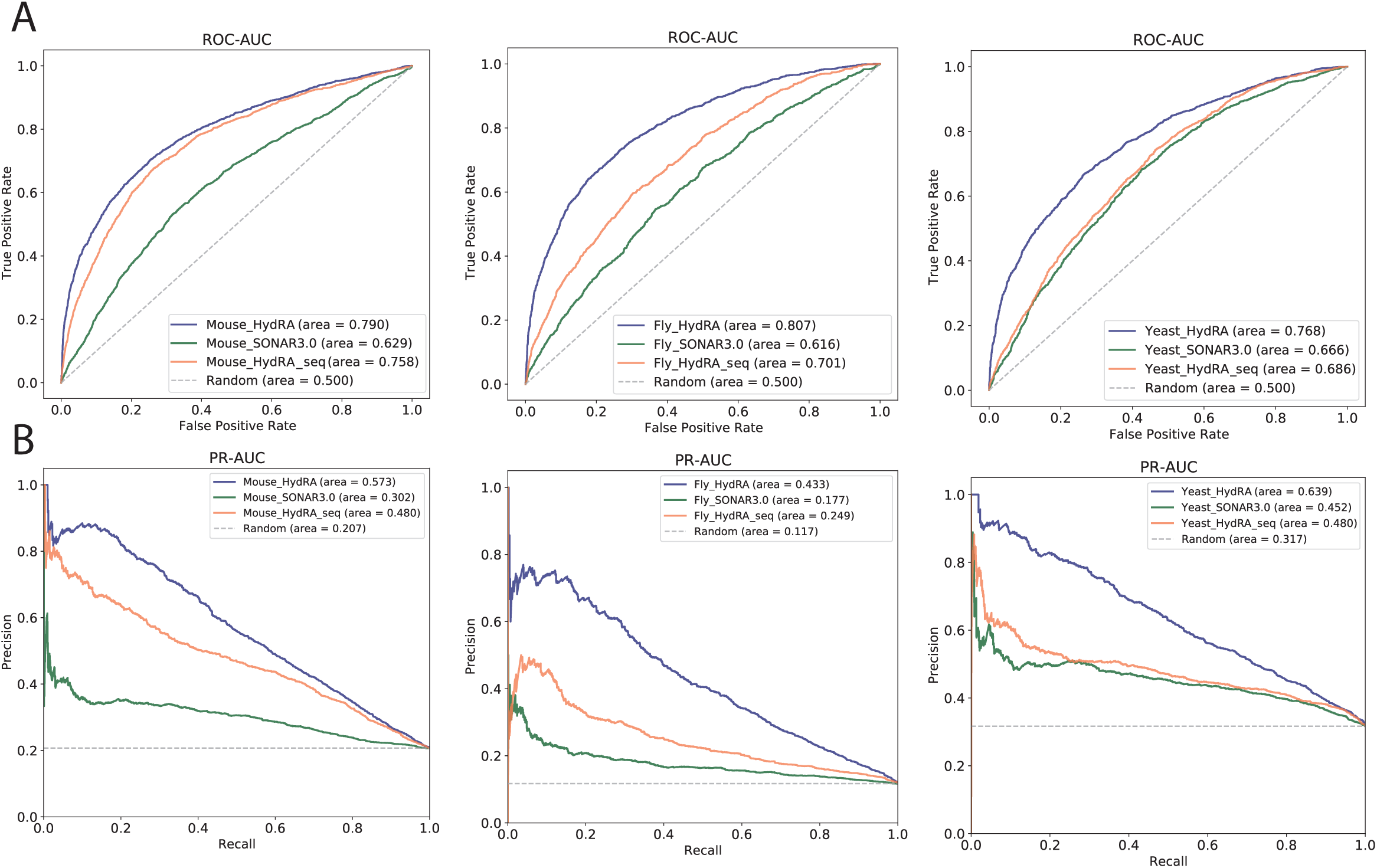
HydRA model trained with human RBPs displays decent predictive power on RBPs from other organisms. **A-B.** Plot showing the ROC-AUC (A) and Precision-Recall (B) performance of HydRA and its sub-classifiers (SONAR3.0 and HydRA-seq) on RBPs from mouse, fly and yeast respectively. To be noticed, the HydRA model and its sub-classifiers are trained with human proteome only. The theoretical random performance is shown as a dashed line.

**Figure S5: Related to Figure 4.**
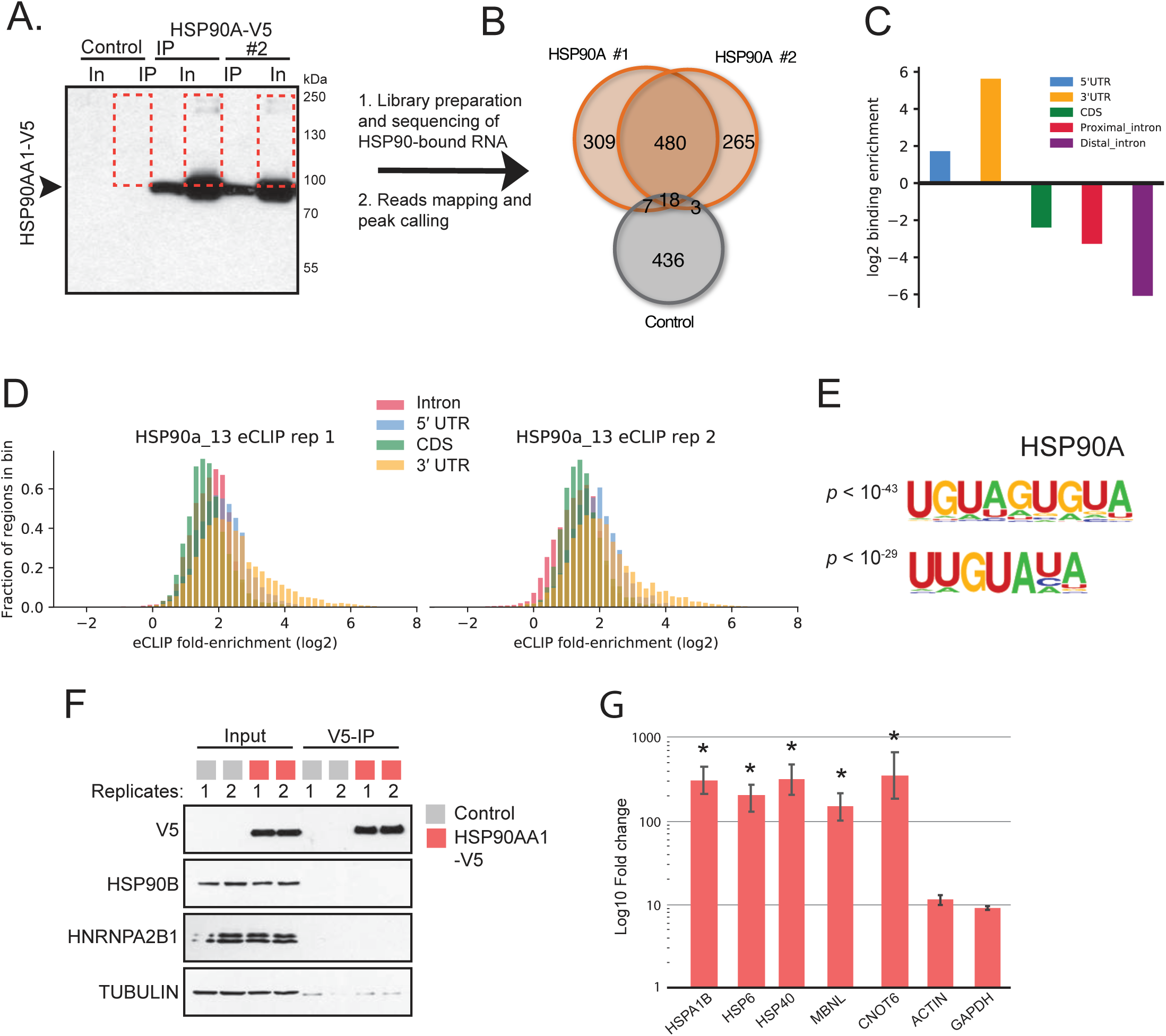
eCLIP analysis discovers HSP90 as an RBP that binds mRNA encoding other heat-shock proteins, and YWHA (14-3-3) family that binds RNAs with similar binding preference across the transcriptome. **A.** Western blot showing immunoprecipitation of HSP90-V5 in HEK293 cells transiently transfected with plasmids expressing V5-tagged HSP90 but not in cells transfected with empty vector (Control). The orange dotted box shows the region of the membrane that excised and RNA isolated for library preparation. **B.** eCLIP-seq reads were mapped to the human genome (hg19) to evaluate RNA expression. Overlap of significant binding peaks on RNAs for each of the replicate HSP90-V5 eCLIP-seq libraries and control library is shown. **C.** Binding preference of HSP90A on different genomic regions measured by log2-transformed region enrichment *E_region_* (see Methods). The following genomic regions of a transcript are shown: 5’UTR (blue), 3’UTR (yellow), coding sequence (CDS, green), proximal introns (red), and distal intron (purple). **D.** Histogram of region-based fold enrichment of sequencing reads for HSP90 (compared to its paired SMInput). 3’UTRs of genes show a higher enrichment of eCLIP signals than the other regions of genes. **E.** Motif logos with corresponding *p* value generated from significant eCLIP peaks of HSP90A. Top two motifs (ranked by *p value*) are shown for each protein. **F.** Western blot analysis to show successful immunoprecipitation of HSP90-V5 for RNA immunoprecipitation. HEK293 cells were transiently transfected with plasmids expressing empty vector (Control) or HSP90-V5. Endogenous HSP90B and HNRNPA2B1 serve as negative controls. TUBULIN serves as a loading control. Experiments were performed in biological duplicates. **G.** RNA immunoprecipitation was performed using a V5 antibody as validated in (G). The relative fold change enrichement of mRNAs by HSP90 was determined by qPCR. Values are fold change ± s.d. for biological duplicates. Student’s t-test was performed between HSP90 and the negative control. Asterisk indicates p < 0.05 and p-values are as follows: HSPAB, 0.00035; HSP6, 0.00364; DNABJ, 0.00114; CNOT6, 0.04792; MBNL1, 0.04532; ACTIN, 0.11138; GAPDH, 0.19121.

**Figure S6: Related to Figure 4.**
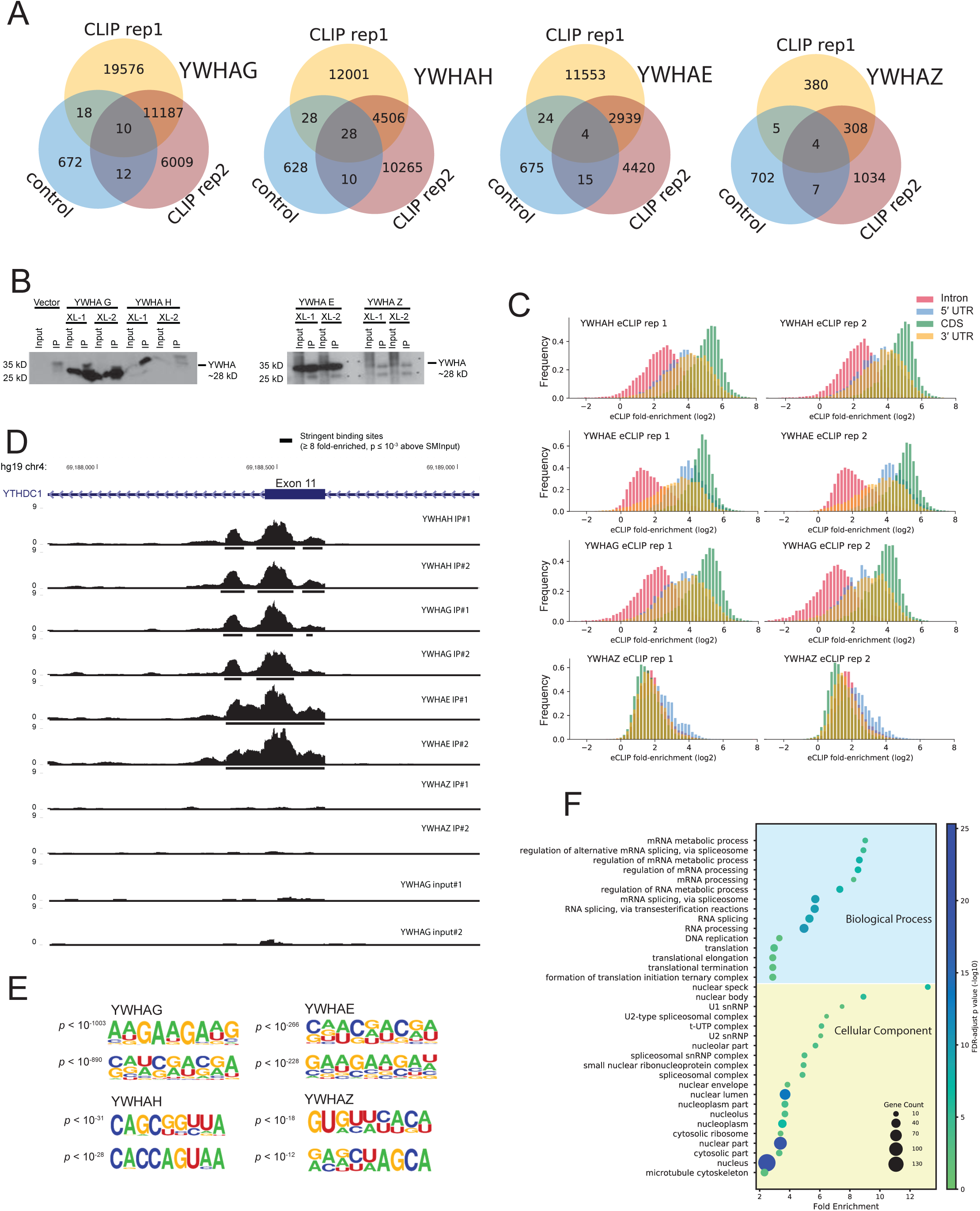
eCLIP analysis discovers that YWHA (14-3-3) family binds RNAs with similar binding preference across the transcriptome. **A.** Overlap of significant binding peaks for each replicate eCLIP-seq libraries and control library for YWHAG, YWHAH, YWHAE and YWHAZ. **B.** Western blot showing immunoprecipitation of YWHAG-V5, YWHAH-V5, YWHAE-V5, YWHAZ-V5, MACROD1-V5 and PIM1-V5 respectively in HEK293 cells transiently transfected with plasmids expressing V5-tagged YWHAG, YWHAH, YWHAE, YWHAZ, MACROD1 or PIM1 but not in cells transfected with empty vector (Control). Re-immunoprecipitation results after immunoprecipitation are also shown for MACROD1 and PIM1 proteins. **C.** Histogram of region-based fold enrichment of sequencing reads for YWHA family members YWHAH, YWHAE, YWHAG and YWHAZ (each compared to its paired SMInput), showing sequencing reads from YWHAH/E/G eCLIP libraries are all enriched in CDS regions, while those from YWHAZ’s are not. **D.** An example of genome browser shots of eCLIP peaks on YTHDC1 gene of YWHAH, YWHAG, YWHAE and YWHAZ eCLIP libraries, showing an shared binding sites of YWHAH and YWHAE with YWHAG, but not with YWHAZ and the SMInput libraries of YWHAG (control). Both the two biological replicates for each proteins are shown. **E.** Motif logos with corresponding *p* value generated from significant eCLIP peaks of YWHAG, YWHAE, YWHAH and YWHAZ. Top two motifs (ranked by *p value*) are shown for each protein. **F.** GO enrichment of the common RNA targets shared by YWHAH, YWHAG and YWHAE. Enriched terms (FDR p < 0.05) with fold-change greater than 2 and with more than 5 observed gene members are shown. To be more informative, only the GO terms from the 7th level in the hierarchy of each GO category are presented.

**Figure S7: Related to Figure 5.**
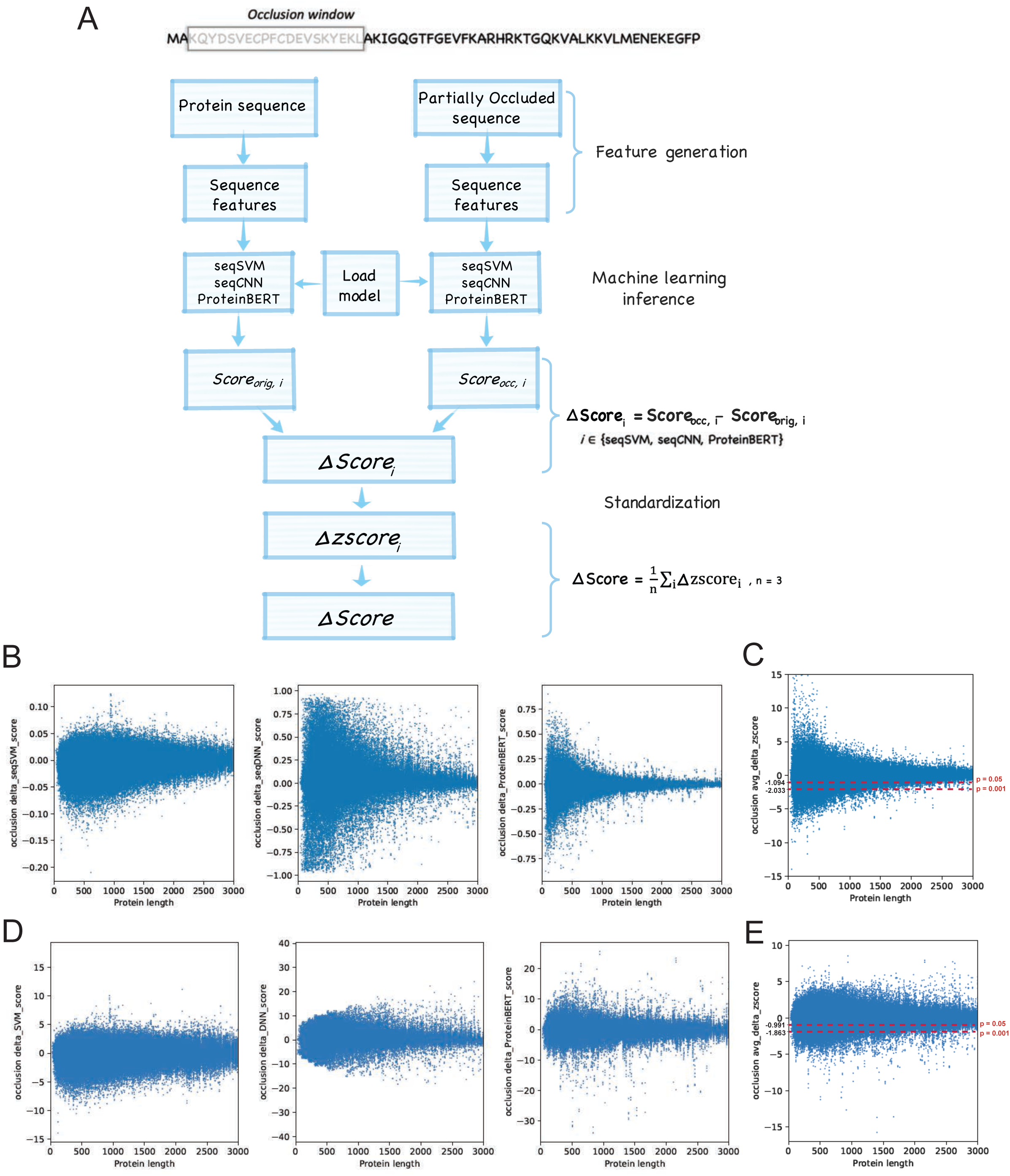
Occlusion map enables model interpretation of HydRA Sequence Classifier and novel RNA-binding domain discovery. A. The workflow that calculating the occlusion score (ΔScore) for each occlusion window along the protein sequence. For simpilicity, occlusion window index *j* was omitted in the equations. B. Scatter plots showing the Δscore_model_ distribution for proteins in different length, where model ∈ {seqSVM, seqCNN, ProteinBERT-RBP}. C. Scatter plots showing the final Δscore distribution from for proteins in different length where no adjustment was done about protein length. D. Scatter plots showing the Δzscore_model_ distribution for proteins in different length, where model ∈ {seqSVM, seqCNN, ProteinBERT}. The standardization step was done within groups of proteins with similar length. E. Scatter plots showing the final Δscore distribution from for proteins in different length where the standardization step was done within groups of proteins with similar length.

**Figure S8: Related to Figure 5.**
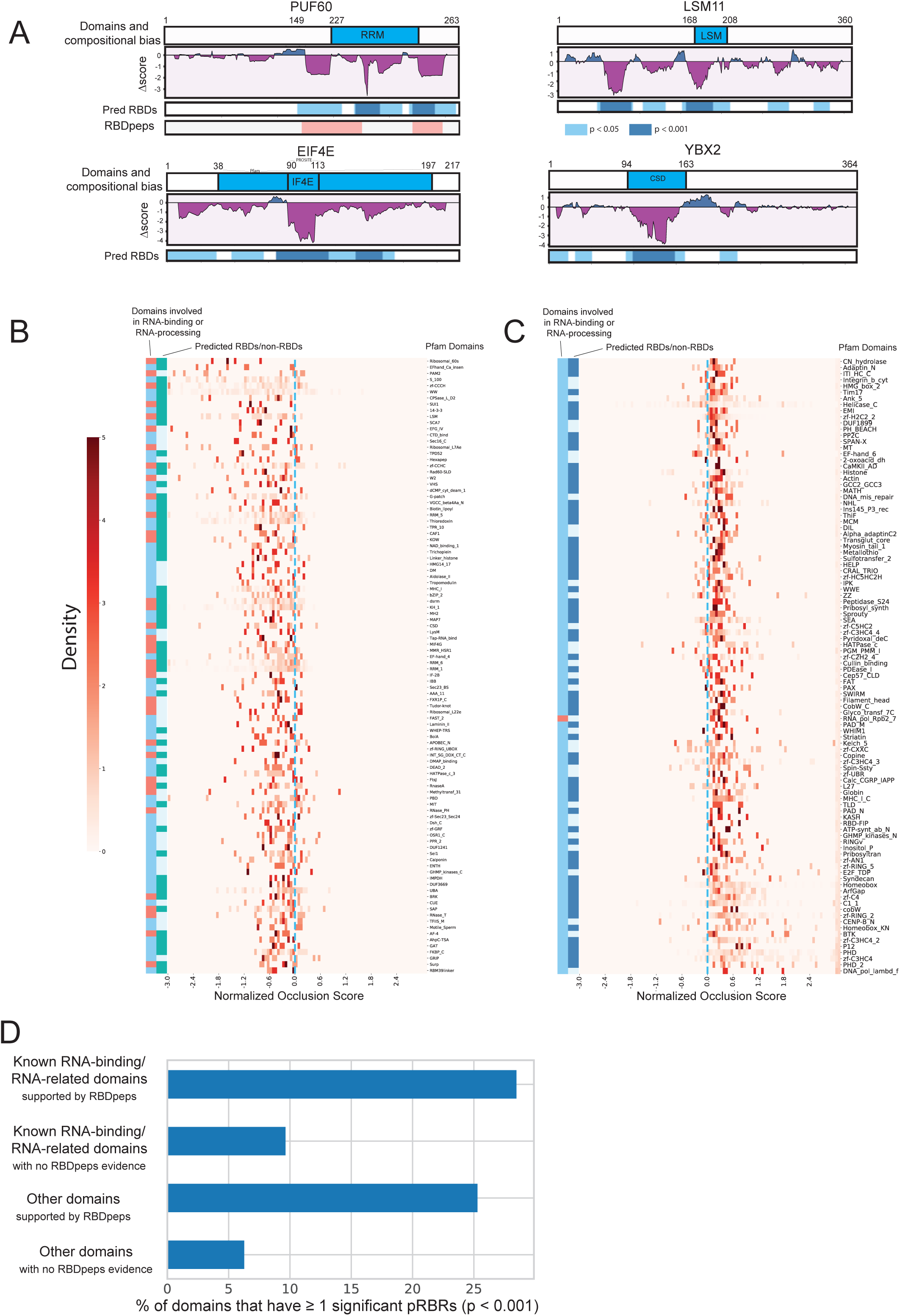
Known and candidate RNA-binding domain discovery with Occlusion Mapping. **A**. Occlusion Maps of several known RBPs (including PUF60, LSM11, EIF4E and YBX2). The coordinates of protein domains, compositional bias and RBDpeps are shown. Protein regions that have *Δscore* < 0 are colored with purple, while those with *Δscore* > 0 are colored with blue in the middle panel. The regions with significant p values are also marked in the bottom track: skyblue for p<0.05 and dark blue for p<0.001. **B, C**. The heatmap of top 100 and bottom 100 domains of Figure 5C. Domains are sorted by their average occlusion scores from small to large. Protein domains that are already known for RNA-binding or involvement in RNA processing are marked in red in the first annotation bar (on the right side) of the heatmap. The second annotation bar shows the predicted RBD (mean < 0 and FDR adjusted p-value < 0.05, one-sample t-test) in green and predicted non-RBD (mean > 0 and FDR adjusted p-value < 0.05) in dark blue. **D.** We grouped the protein domains in known RBPs to four categories: RNA-binding domains (RBDs) that are supported by RNA-binding peptide studies, RBDs that lack support from current RNA-binding peptide studies, other domains with and without evidence from RNA-binding peptide studies. The bar plot represents the percentage of the domains within each category that have ≥ 1 occluders with significant negative *Δscores* (p-value < 0.001).

**Figure S9: Related to Figure 6.**
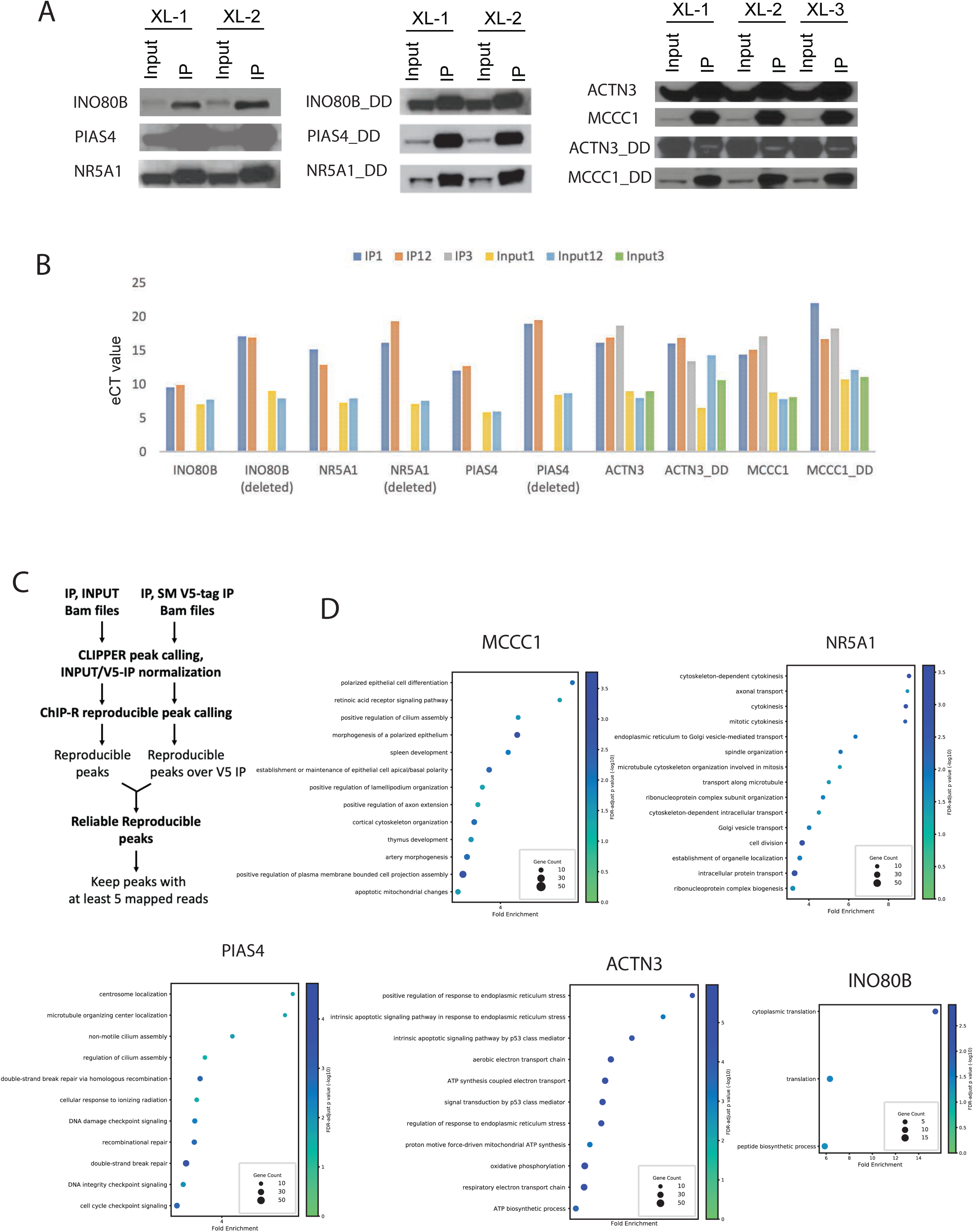
Novel RBPs and functional regions related to RNA-binding were predicted using HydRA and experimentally validated. **A.** Western blot showing immunoprecipitation of V5 tagged candidate RBPs and their corresponding truncated proteins with predicted functional domains deleted in HEK293 cells. U2AF1 is a positive control. **B.** eCT values from the qPCR experiments of eCLIP samples for full-length and truncated candidate RBPs. **C.** The workflow to call reproducible peaks across samples with background signals from V5 tag removed. **D.** Dot plot showing the biological process GO terms enriched in the reliable reproducible protein-coding targets of the candidate RBPs. Statistically significant terms (*FDR p < 0.01*) are shown.

## RESOURCE AVAILABILITY

### Lead contact

Further information and requests for resources and reagents should be directed to and will be fulfilled by the lead contact, Gene Yeo.

### Materials availability

Materials generated by the authors in this study will be distributed upon request.

### Data and code availability

The HydRA software and Occlusion Map are open source and available at https://github.com/YeoLab/HydRA. The repository also includes trained model weights for human RBPs classification.

## STAR Methods

### METHOD DETAILS

#### Mentha-BioPlex network

We obtained a comprehensive protein-protein interaction (PPI) network, i.e. Mentha-BioPlex, by combining BioPlex2.0 ^50^ a high-quality PPI dataset generated by robust affinity purification– mass spectrometry, with a meta-database Mentha ^18^ that collects experimental evidence from different PPI database such as MINT ^19^, IntAct ^21^, DIP ^51^, MatrixDB ^52^ and BioGRID ^20^. Each interaction in the network was assigned a reliability score taking into account all the aggregated experimental evidence, according to the formula defined in MINT database and Mentha:

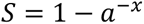

***a*** is a constant. It determines the growth rate of the curve. To keep the scores in a convenient range, we chose ***a***=1.4 as Mentha did. ***x*** is calculated by adding up all the experimental evidence considering type and size of the experiments and the number of publications that support the interaction:

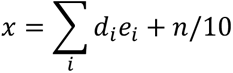

***i*** is the index of all experimental evidence supporting the interaction, while ***d*** takes into consideration the experiment size (d=0.5 for large scale experiment reporting more than 50 interactions, otherwise d=1) and ***e*** reflects the type of information the experiment provides (e=1.0 for evidence of direct interaction and e=0.5 for evidence that only support and association). Additionally, ***x*** also takes into account the number (***n***) of experiments (publications) that support this interaction.

We then presented this network, consisting of only experimentally demonstrated physical protein-protein interactions, as an undirected graph with each protein as the node and the interaction between proteins as the edge. In total, this network contains 19,066 nodes and 309,653 edges.

### The functional protein association network

To supplement current PPI network with extra extrinsic association information of proteins, we introduced functional protein association data into our modelling. We take STRING ^53^ as our functional protein association resource in this study. STRING is a database consisting of protein-protein association data from several sources, i.e. genomic context predictions, high-throughput lab experiments, conserved co-expression, automated text-mining and other PPI databases. To construct the functional protein association network, we kept all the STRING interactions that has at least one protein existing in Mentha-BioPlex network, while the experimentally demonstrated physical protein-protein interactions from STRING are removed to avoid redundancies.

### Protein dataset and Known RBPs collection

On one hand, all the proteins in Mentha-BioPlex network constituted our protein dataset and the corresponding protein sequences were retrieved from Uniprot database. On the other hand, we collected currently known RBPs from published studies ^1,30,40,41, 54–58^, and merged them with proteins in the GO term of “RNA-binding” (GO:0003723) and its descendent terms from AmiGO 2 database (version released in July 2016) ^59,60^, where we removed those proteins assigned by automated methods without curatorial judgement in order to guarantee the high quality. After that, each protein in our protein dataset was assigned a class label, viz. known RBP or not known for RBP (we referred to it as non-RBP for simplicity), based on this RBP list. In all, our dataset contained 18,951 UniProt protein IDs with sequences retrieved, among which 3,762 are known RBPs (also referred to as annotated RBPs).

### Network properties’ effects on SONAR

We studied the effects of PPI network’s properties on the performance of SONAR, such as network density (also referred to as completeness) and the reliability of PPI edges. In this analysis, we first randomly removed certain numbers of edges from the original network that led to a couple of induced networks with retained edges in percentage of 20%, 40%, 60% and 80% respectively from the whole network. For each percentage, we generated 10 different networks as replicates. SONAR was then run with each of these induced networks and the performance was evaluated by ROC-AUC analysis with the output from SONAR software which was obtained by a cross-validation-like (10-fold) approach ^16^. To test if the reliability of PPI edges in the network matters, we also generated a group of induced networks in the same way as above but from a random network that came from an edges-shuffle operation with the original network. ROC-AUC analysis was also done to evaluate the performance of SONAR with these induced networks.

On the other hand, to determine if the degree of reliability of the experimental PPI data matters, we sort the edges in the network based on their corresponding reliability scores (from high to low) and generated a couple of subnetworks from the original network by keeping the top 20%, 30%, 40%, 60% and 80% edges. Similar to the analysis of network density, those induced subnetworks were then run with SONAR and the performance of SONAR was evaluated by ROC-AUC analysis.

### Rationale for HydRA

HydRA is a classification model (also called classifier) aimed at recognizing the RNA-binding capacity of a given protein. It’s the revised version of SONAR ^12^ by introducing new protein features and new machine learning techniques into RBP prediction. HydRA consists of two main components: 1) the first component, referred to as extrinsic classifier, utilizes information describing the extrinsic context of proteins to predict RNA-binding potential from experimental protein interaction data and predicted protein association data (i.e. STRING ^53^); 2) the second component, referred to as HydRA-seq, measures proteins’ RNA-binding potential from their intrinsic properties with their amino acid sequence -based information. Specifically, the second component comprises three sub-classifiers, including a SVM model (seqSVM), a convolutional neural network (seqCNN) model and an attention-based model following Transformer architecture (named ProteinBERT-RBP). The outputs of all the classifiers from the two main components are then combined with a simple probabilistic model to give the final prediction of the input proteins.

### Dataset for modeling

Under the criterion where the class ratio (i.e. known RBPs versus other proteins) in each set keeps same, we first randomly took out 20% of the data in our protein dataset as the hold-out test set, which is only used for the final evaluation of the final RBP classifier and the comparison among different RBP classifiers/software. The rest 80% of data constitutes the training/validation set that is used to train and select model, select features and tune the hyper-parameters of the RBP classifiers. In the evaluation and comparisons of classifiers that utilize proteins sequences, we discarded the proteins in the test dataset whose sequences are highly similar to any protein’s sequence (>90% similarity and >90% coverage) in the training/validation dataset. The similarity between protein sequences was calculated by cd-hit program ^61^.

### Protein interaction and association based classifier

A classifier exploiting the data from physical protein-protein interaction network (i.e. Mentha-BioPlex) and functional protein association network (i.e. STRING), which described the extrinsic context of the protein, was constructed using support vector machine (SVM) model with radial basis function as its kernel. We referred to this classifier as Protein Interaction and Association based (PIA) classifier (also referred to as SONAR3.0). The feature extraction and classifier construction process are modified from that of SONAR.

Firstly, we extracted the local network around the given protein and split the local network into different neighborhoods (level-1, level-2, etc.) according to the length of the path from the protein to the given protein (more details can be found in SONAR (Brannan & Jin & Huelga et al. 2016)). Specifically, we found all the paths with length *k* that start from the given protein and collect all the end nodes of these paths into level-*k* neighborhoods. Notably, a protein can be in both level-1 and level-2 neighborhoods if there are paths in length 1 and 2 between this protein and the given protein. In addition, a protein node can appear more than once in the level-*k* neighborhoods if there are more than one path of length *k* between this protein and the given protein.

Secondly, the proportion of nodes that are annotated RBPs in each neighborhood represents the feature of that neighborhood. In details, for a given protein, its feature *F_k_* of the *k*-th level neighborhood is calculated as

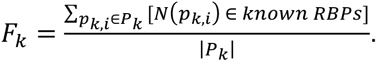

*P_k_* is the set of paths with length = *k* in the network that start from the given protein, *p_k,i_* denotes a specific path in *P_k_* while *i* is the index of the path. *N(p)* denotes the end node of path p. Here […] are the Iverson brackets. [*P*] is defined to be 1 if *P* is true, and 0 if it is false.

The first three levels of neighborhoods in physical PPI network and the first level neighborhood in functional protein association network are considered in this classifier. Different from SONAR, we also added an indicator feature for each local network (i.e. physical PPI local network and functional protein association local network) to penalize those proteins with few level 1 neighbors, whose PPI information is not adequate for confident predictions. We set the indicator feature as 1 if the query protein has no less than 1 level-1 neighbors, otherwise the feature was set as 0. By optimal threshold searching, we set 1 = 5.

Lastly, we took the six network-based features of each given protein as the input of SVM model. Cross-validation (10-fold) method was used to search for the optimal hyper-parameters and threshold (such as the number of PPI neighborhoods included and the indicator threshold 1) of this model within our training/validation set. Additionally, to handle the class imbalance issue in our training process, we chose oversampling out of a couple of widely used approaches, where we oversampled the number of our RBP training samples by equally replicating current RBP samples in the training set.

### Sequence-based SVM classifier

In line with many previous studies on sequence-based RBP prediction, a support vector machine (SVM) model with radial basis function (RBF) kernel was used to construct one of the sequence-based RBP sub-classifiers from protein sequence. We referred to this classifier as seqSVM.

Firstly, protein sequences were preprocessed to generate three categories of protein features:

1. *k-mer*: all the possible substrings of length *k* (*k=3, 4*) that are contained in a protein’s amino acid sequence were counted. The counts of all k-mers were subsequently encoded into an integer vector. To alleviate the computational cost, for each protein, we only look at the 1000 most frequent *k-mers* that appeared in the known RNA-binding proteins.
2. Amino acid composition (AAC): We calculated AAC for each protein with *propy* ^62^ package. A vector of 20 descriptors per protein was gained.
3. *k-mer* of predicted secondary structure (*SS-kmer*): Secondary structure type of each amino acid in the protein was predicted by SPIDER2 software ^63^. The induced sequence of secondary structures of the protein was then used to generate the k-mer (*k=11, 15*) feature vector in the same way described in (1) except that we look at the 1500 most frequent *SS-kmers* that appeared in the known RBPs.

Secondly, feature selection was done in order to reduce the dimension of feature vectors. Chi-square test was used to filter out unimportant features within each feature category respectively via their *p* values controlled for false discovery rate (FDR) at level 0.01. For *k-mer* and *SS-kmer*, features were further selected by linear SVM model by keeping the features the absolute value of whose weights are larger than the average of those of all input features.

Thirdly, the seqSVM classifier was built with the features selected from last step utilizing SVM model and RBF kernel. We only use proteins shorter than 1500 amino acids in our training step to alleviate the involvement of unrelated information carried by proteins, particularly those long ones.

We used the training/validation set with cross-validation approach to search for optimal values for *k* (in *k-mer* and *SS-kmer*) and parameters or hyper-parameters used in feature selection pipeline and SVM model construction. The same oversampling approach as mentioned in SONAR3.0 construction was employed to overcome the class imbalance issue in the training process.

### Convolutional neural network for sequence-based RBP classification

Parallel to seqSVM, we also created a feed forward convolutional neural network (CNN) to recognize RBPs by automatically learning the pattern underlying the protein sequence with convolutional layers. We referred to this CNN model as seqCNN. The seqCNN model consisted of 8 different hidden layers of neuron-like computational units that are stacked up in a certain order (shown in **Figure 2A**). The first layer was an embedding layer that mapped amino acids to dense vectors encoded with biophysical and biochemical properties of the amino acids using a pre-trained weight matrix called ProtVec ^23^. This layer not only enriches protein sequence information with the knowledge from available database, but also alleviates the difficulties on deep learning caused by sparse one-hot encoded matrix of the protein sequence. Following this layer were two 1D convolutional layers (kernel size = 5) with a pooling layer for each to extract patterns from sub-sequences and condense the information for downstream layers. The output was then followed by a global pooling layer for further condensation. Next, the condensed information was conveyed to last 2 hidden layers, which are regular fully-connected neural network layers (also called dense layers), for more processing. An output layer in the form of fully connected neurons after the last hidden layers is used to encode all learned protein information into the form of probability (i.e. the classification score), showing how likely the input protein is a RNA-binding protein. The binary-cross-entropy loss was used to measure the agreement between the output predictions and the true class labels the on the training set. Besides, regularization methods such as dropout and batch normalization are applied to avoid overfitting. We choose this model structure from a broad range of CNN topologies (i.e. the number of CNN layers, the number of kernels in each layer and the) and alternative CNN architectures, such as residual convolutional neural network.

This model was trained with backpropagation algorithm that adjusts and optimizes the weight matrix in each layer during the training process to minimize the binary-cross-entropy loss on the training set. Specifically, Adam optimization was used in the process. To prevent the training process from getting stuck in undesired local optima, we pre-trained the hidden layers with auto-encoder technique ^64^ where the convolutional layers and fully-connected layers were pre-trained separately. Similar to seq SVM, only proteins with length shorter than 1500 amino acids are used to train the model and all the optimal hyper-parameters of this model were obtained using cross-validation within our training/validation set. The seqCNN classifier was implemented with TensorFlow kernel ^65^ utilizing its high-level API, Keras ^66^. The class imbalance issue in the training process was handled by adjusting the class weight argument in Keras model.

### Attention-based protein language model for RBP classification

ProteinBERT is an attention-based Transformer/BERT architecture specifically designed for protein classification problems ^15^. It originally takes protein sequence and protein GO annotations as input. The key component in this architecture is a the transformer block with two parallel neural networks: (1) one takes the representation matrix from the protein sequence (denoted by local representations) and uses convolutional layers with skip connections and layer normalizations to extract and compress location-wise signals followed by (location-wise) fully connected layer; (2) the other consists of two fully-connected layers with skip connections and layer normalizations and takes a global representation vector encoded from the GO annotations of the protein as input. In each transformer block, information between local and global representations flows to each other via two attention layers respectively allowing the local and global signals to guide the learning process of each other easily ^15^. We employed the same ProteinBERT architecture as proposed in their original paper which includes 6 transformer blocks with 4 global attention heads in each block.

This ProteinBERT model was pretrained following the original paper via a self-supervised training strategy with ∼106M proteins and their corresponding GO annotations from UniRef90 ^67^. We next fine-tune the model with our training set with RNA-binding annotations (mentioned above) allowing the model to exclusively focus on the supervised learning of RNA-binding related sequence patterns and to classify RBP and non-RBPs. In the fine-tuning, all layers of the pretrained model were first frozen except a newly added fully-connected layer in the end of the model, which was used to connect the pretrained model to RBP classification output layer. Similar to seqCNN, binary-cross-entropy loss was used to measure the agreement between the output predictions and the true class labels and Adam was used in the optimization process. The model was trained for up to 40 epochs. Next, we unfroze all the layers and trained the model for up to 40 additional epochs. Lastly, we did a 1 final epoch of training with a larger sequence length following the original paper ^15^. We evaluate the model using the same strategy of the classifiers above with training and test set while 10% of the training set was used as “validation set”. The model was fine-tuned using Adam optimizer for the backpropagation, and we reduced the learning rate on plateau and used early-stopping base on the “validation set” to avoid overtraining. This ProteinBERT model for RBP classification (referred to as ProteinBERT-RBP) was implemented using Keras ^66^ with Tensorflow kernel ^65^.

### Graph neural network modelling on protein structures

The protein structure backbone is represented as graphs G = (V, E), where each node *ν* ∈ V is an amino acid connected by edge 4 ∈ E to its amino acid neighbors in the 3D space whose corresponding Ca atoms are less than 10Å distant from the C_*α*_ atom of *ν* or within the k-neareast neighbor set of the C_*α*_ atom of *ν*. To obtain the optimal features to represent each amino acid and their spatial relationship to their neighbors, we select node features from various types of amino acid features including (1) one-hot representation, (2) main chain torsion angles (Phi and Psi), (3) the structural features encoded by FoldSeek^68^, and the latent representations learned by large pre-trained protein language models, such as (4) DeepFRI^26^, (5) ProteinBERT^69^, and (6) HydRA’s ProteinBERT-RBP. Similarly, we select edge features from an edge attribute pool including (1) using the same attributes for each edge (i.e. no edge features) (2) distance between C_*α*_ atom of the two amino acids, (3) the structural features encoded by FoldSeek between the two amino acids, (4) the average confidence score (pLDDT) from AlphaFold2 prediction of the two amino acids. The feature selection was done using the forementioned training set and validation set where the models with different combination of the node and edge features were trained on the protein structures with RBP labels in the training set and evaluated on the validation set. The optimal graph neural network architecture was also selected in this step from the widely used architectures, e.g. graph convolutional network (GCN), graph attention network (GAT) and PointTransformer ^70^. The optimal model was selected and denoted as strucGNN1 which employs PointTransformer as the main architecture. The local presentation output for amino acid (node *ν_i_*) by the last transformer block in ProteinBERT-RBP model was selected as the node features in strucGNN1, defined as *x*1*_i_* = ProteinBERT_RBP(*ν_i_*), while no edge feature was used as PointTransformer architecture does not support edge feature input.

To update the node features of each amino acid according to the protein structure graph, strucGNN1 use 3 PointTransformer blocks. Each PointTransformer block firstly down-samples the nodes in the graph using farthest point sampling and then aggregates information from the neighboring nodes for each node using self-attention mechanism ^70^. In details, the node-wise aggregation operation in k-th PointTransformer block is defined as:

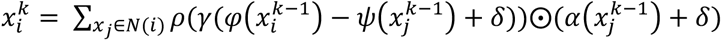

where *N*(*i*)denotes the neighbors of *ν_i_*, and *ν_i_* is denoted as *i* here for simplicity. and 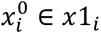. As defined in the original paper for PointTransformer ^70^, *φ*, *ψ*, and *α* are pointwise feature transformations using fully-connected layers, *δ* is a position encoding function used to encode the relative position between two nodes. *δ* is defined as *δ* = *θ*(*p_i_* − *p_j_*, where *p_i_* and *p_j_* are the 3D coordinates of the C*α* atom for each node and the encoding function θ is two linear layers with one ReLU nonlinearity. *ρ* is a normalization function *softmax*, while the mapping function γ is also two linear layers and one ReLU nonlinearity. In this way, each block applies self-attention locally, within a local neighborhood around each amino acid.

The global features of the protein are then calculated by taking the channel-wise average across the node dimension output by the PointTransformer blocks using a global mean pool layer, followed by two fully-connected layers (with 64 neurons each) with ReLU non-linear activation function to further process the global features. The final output is computed as a linear mapping of the fully-connected layer’s output followed by Sigmoid transformation. Binary-cross-entropy loss was used to measure the agreement between the output predictions and the true class labels and Adam was used in the optimization process. The strucGNN1 were implemented using PyTorch^71^ and PyTorch-Geometric ^72^.

### Finetuning pretrained Graph neural network on RNA-binding protein structures

To exploit the recent advancement in the structure-based protein pretraining models, we adjusted and finetuned Masked Inverse Folding (MIF) model for RBP classification task and referred this new model as strucGNN-MIF. MIF is a pretrained graph neural network model which achieved state-of-the-art performance in many protein prediction tasks ^73^, MIF was pre-trained with the CATH4.2 database^74^, and is built on structured graph neural network architecture with the pretraining task of reconstructing a corrupted protein sequence conditioned on its backbone structure.

Similar to the graph construction in strucGNN1 and strucGNN2, the protein structure backbone is represented as graphs G = (V, E) with each amino acid as a node *ν* ∈ V. But instead of taking the other amino acids with the distance less than 10 Å to the given amino acid as neighbors, each given amino acid node is connected by edge 4 ∈ E to its k-nearest amino-acid neighbors in the structure.

To adjust MIF to binary RBP classification task, we took the encoder layers of MIF and connected it to a global mean pool layer to average node features across the node dimension. The pooled features were then fed to three fully-connected layers with ReLU non-linear activation before connecting to the output layer with sigmoid activation function.

We finetuned this model with our RBP training set and evaluate the performance on the validation and test set. Similar to strucGNN1 and strucGNN2, binary-cross-entropy loss was used to measure the agreement between the output predictions and the true class labels. Adam was used to adjusts and optimizes the weight matrix in each layer and to minimize the binary-cross-entropy loss on the training set.

### Meta ensembling of classifiers

For simplicity, we denoted the aforementioned three classifiers (i.e. SONAR3.0, seqSVM and seqDNN) as primary classifiers. A simple probability-based ensemble approach was then derived to integrate predictions from these primary classifiers. This approach consists of three steps:

1. We collected the classification scores generated by each primary classifier during the cross-validation step and took these scores and corresponding class labels as reference lists *S* and *L*:

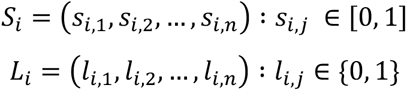

where *i* is the index of the primary classifier, *j* is the index of the proteins and *n* represents the total number of protein scores collected from the cross-validation step.
2. For a query protein, given a classification score *x* from a primary classifier, we calculated the probability of this protein being a false discovery (i.e. false discovery rate) for this primary classifier, based on *x* and the reference lists *S* and *L*:

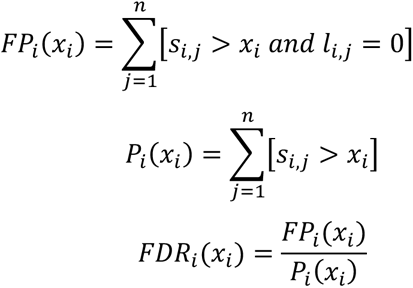

where *FDR* represents false discovery rate, *FP* represents the number of false positives and *P* represents the number of all the positive prediction. Here […] are the Iverson brackets. [*P*] is defined to be 1 if *P* is true, and 0 if it is false.
3. For this query protein, we then got its probability of being a false discovery for all these primary classifiers as follows:

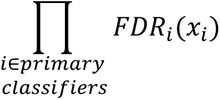
4. Thus, the probability of this query protein not to be a false discovery for at least one of the primary classifiers was got as follows. This is taken as the output score of this ensemble approaches.

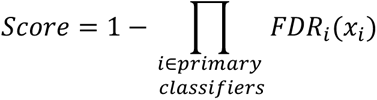

Besides constructing the final RBP classifier (i.e. HydRA), this ensemble approach was also used to integrate the results from intrinsic classifiers (i.e. seqSVM and seqDNN)

This ensemble approach is chosen from a couple of other classifier ensemble methods, including other variant of this probability-based approach (i.e. the false positive rate based version) and machine learning based approaches such as SVM and logistic regression.

### Software comparisons

We applied other publicly available RBP classifiers, i.e. RNApred ^7^, SPOT-seq ^8^, catRAPID signature ^9^ and RBPPred ^10^ to predict all the proteins in our test dataset. The predicted scores and class labels were then collected to calculate model evaluation metrics such as accuracy, sensitivity, specificity, precision, ROC-AUC, F1 score, Matthews Correlation Coefficient and balanced accuracy.

### RNA-binding domain sequences and RNA-binding peptide

Amino acid sequences corresponding to different protein domains (i.e. KH, RRM, DEAD, CSD, LSM, BTB, SH2, Ig, UBA and 7 transmembrane domains)) are downloaded from Pfam database. RNA-binding peptides are collected from five recent studies of four technologies that experimentally identify the regions that interact with RNA molecules in RNA-binding proteins: RBDmap ^6,44^, pCLAP ^45^, RBR-ID ^75^ and RNP^xl^ ^46^.

### Protein domain annotation

The domain annotations in each protein are obtained by homology search using hmmscan (HMMER 3.1b1) against Pfam-A.hmm from Pfam database (v27.0). Overlapped and redundant domain hits are merged using hmmscan-parser.sh (from https://github.com/carden24/Bioinformatics_scripts/blob/master/hmmscan-parser.sh). Only domain annotations with E-value < 0.1 were kept for downstream analysis.

### Occlusion Map

Occlusion map (also referred to as occlusion analysis) aims at interpreting the foci of the machine learning classifier on their target object by detecting the changes in predictions when a part of the object is deliberately “occluded”. This analysis has been widely used in computer vision. Here, we use it to interpret which parts of the protein our intrinsic-feature classifier is “paying attention” to. Specifically, shown in **Figure S7A**, we occluded a subsequence of fixed length *k* (*k*=20 in this study) within the protein sequences by setting a window of consecutive sites to be zero or null (referred to as “occlusion window”). The occluded protein then went through all our feature generation stages and was fed to seqCNN, seqSVM and ProteinBERT-RBP to generate new classification scores (noted as *Score_occ, seqCNN_*, *Score_occ, seqSVM_* and *Score_occ, ProteinBERT_*). We defined the difference between the new classification scores and original scores without occlusion (denoted by *Score_orig, seqSVM_*, *Score_orig, seqDNN_* and *Score_orig, ProteinBERT_*) as Δ *score_model,j_* = *Score_occculed,model,j_* − *Score_original,model_*, where *j* indicates the index of the occluded part and *model* is either seqCNN, seqSVM or ProteinBERT-RBP. In this case, a negative *Δ score* represents a negative effect on categorizing this protein to RBP, indicating the occluded part are considered as important for RNA-binding by our classifier. To integrate occlusion information from seqCNN, seqSVM and ProeinBERT-RBP, Δ *Score_seqDNN_*, Δ *Score_seqSVM_* and Δ *Score_proteinBERT_* were first standardized (i.e. z-score transformed) respectively as 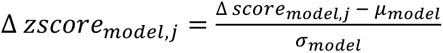, where *μ* and *σ* represents the mean and standard deviation of all Δ scores we have obtained from all the training proteins for specific model (i.e. seqCNN or seqSVM). The normality of Δ *score_seqDNN_*, Δ *score_seqSVM_* and Δ *score_proteinBERT_* populations were confirmed with D’Agostino and Pearson’s test. The integrated Δ *score* was then generated by averaging the standardized Δ *score* from seqCNN and seqSVM: 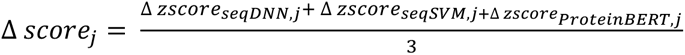. To get a map of *Δ score* for the whole protein, we slid the occlusion window within each protein from N terminal to C terminal and repeated the calculations. P-value was obtained for each occlusion window based on the *Δ score* population which follows a normal distribution. We defined a predicted RNA-binding region (pRBR) as the occlusion window with significantly lower Δ *score* with p-value < 0.05 and a strong predicted RNA-binding region (strong pRBP) with p-value < 0.001. Notably, Δ *score_model_* tends to get smaller as the protein length increases (see **Figures S7B** and **S7C**) so that occlusion windows in short proteins are more likely to be called as pRBR or strong pRBR than those in long proteins. To alleviate this bias, we grouped the proteins based on their protein lengths and do the standardization within each group in the standardization step and this results in a more balanced Δ *score_model_* distribution (**Figures S7D** and **S7E**).

### Criteria of RBP candidacy

After all the aforementioned steps of model selection and optimization, we got the final RBP classifier (i.e. HydRA) and trained it with all the samples in our protein dataset. With this trained classifier, we got a classification score to each protein in the protein dataset. We set a cutoff as 0.8927 for the classification scores by fixing the false positive rate to 10% within the test dataset. Thus, all those proteins with classification scores higher than the cutoff but with negative class labels (i.e. previously not known for RNA-binding) were collected in the list of candidate RBPs.

### Biophysical properties of proteins

Within each RBP candidate, isoelectric points were calculated with ExPASy web server ^76^. A score describing intrinsic disorder was assigned to each amino acid position using IUPred ^77^. Intrinsic disordered region (IDR) or sites were defined as positions with score < 0.4. Complexity of each amino acid within the protein was calculated as Shannon entropy of the neighbor amino acids within a window of size 21 centering the given amino acid’s position. Positions with entropy < 3 were regarded as low complexity region or sites. One-side Mann-Whitney U test was used to statistically examine the difference between different protein populations in the distributions of isoelectric points, amino acid composition and the proportion of disordered and low-complexity regions.

### Antibodies for western blot and immunoprecipitation

The primary antibodies used are as follows: anti-V5 (Proteintech, 66007-1-Ig), anti-V5 (Bethyl A190-120A), anti-His (Proteintech, 66005-1-Ig), anti-His (Invitrogen, MA1-21315), anti-a-Tubulin (Sigma, Clone B-5-1-2), anti-HSP90B (Sigma, SAB4501463) and anti-HNRNPA2B1 (Proteintech, 14813-1-AP)

### Plasmid construction

All RBP mammalian expression constructs were in one of two lentiviral Gateway (Invitrogen) destination vector backbones: (1) pLIX403_V5_mRuby or (2) pLIX403_V5. The pLIX403 inducible lentiviral expression vector was adapted from pLIX_403 (deposited by D. Root; Addgene plasmid no. 41395) to contain TRE-gateway-mRuby and PGK-puro-2A-rtTA upstream of mRuby by Gibson assembly reaction of PCR products (Cloneamp, Takara Bio). RBP open-reading frames (ORFs) were obtained from human Orfeome 8.1 (2016 release) donor plasmids (pDONR223) when available, or amplified (Cloneamp, Takara Bio) from cDNA obtained by SuperScript III (Invitrogen) RT–PCR of HEK293XT cell purified RNA (Direct-zol, Zymogen) and inserted into pDONR223 by Gateway BP Clonase II reactions (Invitrogen). Donor ORFs were inserted in frame upstream of V5 and mRuby fusion cassettes by gateway LR Clonase II reactions (Invitrogen).

Putative RNA binding domains (as determined by occlusion analysis) were removed via site-directed mutagenisis (NEB E0554S) by amplifying entire entry clone plasmids (candidate RBP ORFs within pDONR221 or pDONR223 plasmids) using standard, non-mutagenic forward and reverse primers flanking the deletion region, followed by PCR product ligation. The resulting deletion clones were used for Gateway insertion into V5 or V5-mRuby for stable expression and eCLIP as was done for deletion or full-length expression clones.

### Generation of Cell lines

Stable lines were made in human lenti-X HEK293T cells (HEK293XT, Takara Bio), maintained in DMEM (4.5 g l−1 D-glucose) supplemented with 10% FBS (Gibco) at 37 °C with 5% CO2. Cells were passaged at 70-90% confluency by rinsing cells gently with DPBS without calcium and magnesium (Corning) then dissociating with TrypLE Express Enzyme (Gibco) at a ratio of 1:10. The stable HEK293XT cell lines expressing RBP full-length and domain-deleted RBP candidates and were generated as described below by transducing ∼1 million cells with 8 µg m l−1 polybrene and 1 ml viral supernatant in DMEM + 10% FBS at 37 °C for 24 h, followed by subsequent puromycin resistance selection (2 µg ml−1).

Lentivirus was packaged using HEK293XT cells seeded approximately 24 h before transfection at 30–40% in antibiotic-free DMEM and incubation at 37 °C and 5% CO2 to 70– 90% confluency along with the Lipofectamine 3000 reagent (ThermoFisher Scientific) following the manufactures protocol (https://tools.thermofisher.com/content/sfs/manuals/lipofectamine3000_protocol.pdf). DNA ratios used for transfection is as follows: 10:1:10 proportion of lentiviral vector:pMD.2g:psPAX2 packaging plasmids. Six hours following transfection, medium was replaced with fresh DMEM + 10% FBS. At 48 h after medium replacement, virus-containing medium was filtered through a 0.45-μm low-protein binding membrane. Filtered viral supernatant was then used directly for line generation by transducing ∼1 million cells (one well of a six-well dish) with 8 µg ml−1 polybrene and 1 ml viral supernatant in DMEM + 10% FBS at 37 °C for 24 h. After 24 h of viral transduction, cells were split into 2 g l−1 puromycin and selected for 72 h before passaging for storage and downstream validation and experimentation.

### RNA immunoprecipitation and qPCR

HEK293 cells were transfected with plasmids to express V5-tagged HSP90AA1 (pcDNA6-HSP90AA1-V5) or empty vector. 24 hours later, cells were lysed in lysis buffer (50 mM Tris pH 7.4, 100 mM NaCl, 1% NP-40, 0.1% SDS and 0.5% sodium deoxycholate) supplemented with 1 Protease Inhibitor cocktail (Roche) and 80 U of RNAse Inhibitor (Roche). Clarified lysates were pre-cleared with Protein G agarose beads (Life Technologies). Aliquots of the supernatant (equivalent to 5% of supernatant) were saved as input protein and RNA. The remainder of the supernatant was incubated with 2 mg of V5 antibody at 4 C for 4 h. The protein–RNA–antibody complex was precipitated by incubation with Protein G magnetic beads overnight at 4 C. Beads were washed twice with lysis buffer and three times with wash buffer (5 mM Tris pH 7.5, 150 mM NaCl, 0.1% Triton X-100). Ten per cent of the bead slurry was reserved for western blot analysis. The remaining bead slurry was resuspended in TRIzol (Life Technologies), and RNA was extracted as per the manufacturer’s instructions. Input and immunoprecipitated RNA was converted into cDNA and gene expression was measured with qPCR in technical triplicates. RNA immunoprecipitation qPCR studies were performed in biological duplicates.

### eCLIP Library Preparation for YWHAG/H/E/Z and HSP90AA1

The open reading frame of human HSP90AA1 was cloned into pEF5/FRT/V5-DEST (Invitrogen) by Gateway cloning, positioning the gene in frame with a 3’ V5 tag. The open reading frames of human YWHAG, YWHAH, YWHAE, YWHAZ, and HSP90AA1 were cloned into pcDNA6/myc-His B (Invitrogen) by restriction digest, positioning the gene in frame with a 3’ myc-his tag. HEK293T cells transfected with the protein of interest were cultured to confluency in 10 cm dishes. The cells were UV crosslinked (254 nm, 400 mJ/cm2 constant energy) then pelleted and frozen on dry ice. eCLIP procedure was performed as described ^48^. Briefly, cell pellets were lysed in eCLIP lysis buffer and sonicated (BioRuptor). Lysates were treated with RNase I (Ambion) to fragment the RNA and incubated with antibodies against V5-tag or His-tag (both from Proteintech) to immunoprecipitate RBP-RNA complexes. 2% of the lysate:antibody mixture was saved as input sample. The remaining immunoprecipitated (IP) samples were stringently washed, followed by dephosphorylation of RNA by FastAP (ThermoFisher Scientific) and T4 Polynucleotide Kinase (NEB), and ligation of a 3′ RNA adapter with T4 RNA Ligase (NEB). Immunoprecipitates and input samples were resolved by SDS-PAGE, transferred to nitrocellulose membrane, and then the region of membrane corresponding to the molecular weight of the protein of interest up to 75 kDa above it was excised for each immunoprecipitate and input sample. RNA was isolated from the membrane, reverse transcribed with AffinityScript (Agilent), free primers were removed (ExoSap-IT, Affymetrix), and a 5’ DNA adapter was ligated onto the cDNA product with T4 RNA ligase (NEB). Libraries were then amplified with Q5 PCR mix (NEB), size selected using AMPure XP beads (Beckman Coulter, Inc.) and on a 3% agarose gel, and then quantified with a Bioanalyzer instrument (Agilent). Paired-end (50 base pair) sequencing of the eCLIP libraries were performed on Illumina HiSeq 3500.

### eCLIP Library Preparation for INO80B, PIAS4, NR5A1, ACTN3, and MCCC1

All eCLIPs were conducted following induction or transient transfections and IP was conducted using anti-V5 tag (Bethyl A190-120A). eCLIP experiments were performed as previously described in a detailed standard operating procedure6 (6. Van Nostrand, E. L. et al. Robust transcriptome-wide discovery of RNA-binding protein binding sites with enhanced CLIP (eCLIP). Nat. Methods 13, 508–514 (2016).) which is provided as associated documentation with each eCLIP experiment on the ENCODE portal (https://www.encodeproject.org/documents/fa2a3246-6039-46ba-b960-17fe06e7876a/@@download/attachment/CLIP_SOP_v1.0.pdf/). In brief, 20 million cross-linked cells were lysed and sonicated, followed by treatment with RNase I (Thermo Fisher) to fragment RNA. Antibodies were precoupled to species-specific (anti-rabbit IgG) dynabeads (Thermo Fisher), added to lysate and incubated overnight at 4 °C. Before IP washes, 2% of sample was removed to serve as the paired-input sample. For IP samples, high-salt and low-salt washes were performed, after which RNA was dephosphorylated with FastAP (Thermo Fisher) and T4 PNK (NEB) at low pH, and a 3′ RNA adapter was ligated with T4 RNA ligase (NEB). Ten percent of IP and input samples were run on an analytical PAGE Bis-Tris protein gel, transferred to PVDF membrane, blocked in 5% dry milk in TBST, incubated with the same primary antibody used for IP (typically at 1:4,000 dilution), washed, incubated with secondary horseradish peroxidase-conjugated species-specific TrueBlot antibody (Rockland) and visualized with standard enhanced chemiluminescence imaging to validate successful IP. Ninety percent of IP and input samples were run on an analytical PAGE Bis-Tris protein gel and transferred to nitrocellulose membranes, after which the region from the protein size to 75 kDa above protein size was excised from the membrane, treated with proteinase K (NEB) to release RNA and concentrated by column purification (Zymo). Input samples were then dephosphorylated with FastAP (Thermo Fisher) and T4 PNK (NEB) at low pH, and a 3′ RNA adapter was ligated with T4 RNA ligase (NEB) to synchronize with IP samples. Reverse transcription was then performed with AffinityScript (Agilent), followed by ExoSAP-IT (Affymetrix) treatment to remove unincorporated primer. RNA was then degraded by alkaline hydrolysis, and a 3′ DNA adapter was ligated with T4 RNA ligase (NEB). Quantitative PCR was then used to determine the required amplification, followed by PCR with Q5 (NEB) and gel electrophoresis for size selection of the final library. Libraries were sequenced on the HiSeq 2000, 2500 or 4000 platform (Illumina). Each ENCODE eCLIP experiment consisted of IP from two independent biosamples, along with one paired size-matched input (sampled from one of the two IP lysates before IP washes).

### eCLIP Library Preparation for V5 tag control

A plasmid encoding the V5 peptide was transfected in HEK293XT cells using Lipofectamine 3000 in biological duplicates (10 cm plates). After 72hr, cells were UV-crosslinked and subjected to the eCLIP protocol using anti-V5 antibody (Bethyl A190-120A, 10 ug per sample), as previously described. At the SDS-PAGE step, each sample was divided equally between three lanes. Membrane regions corresponding to each possible 75 kDa window, each separated by 25 kDa, were excised and prepared separately for sequencing. Samples were compared against a size-matched input without V5 IP for each size range. We referred to this set of data to as “V5 tag only” samples.

## QUANTIFICATION AND STATISTICAL ANALYSIS

### Evaluating domain-level RNA-binding element prediction

To evaluate HydRA’s capability of recognizing RNA-binding elements on domain level, we ran HydRA-powered Occlusion Map with all the known RBPs and measure the performance of recognizing high-confidence RNA-binding domains (RBDs) in the RBPs over other domains using positive likelihood ratio (discussed below). To obtain the annotation of high-confidence RNA binding domains, we first retrieved the homology-annotated domains (see ***<u>Protein</u> <u>domain</u> annotation*** above) for each RBP and retained the domain hits that exist in the list of RNA-binding and -processing domains from Gerstberger et al. ^1^. Next, those homology-annotated RBDs that were also supported by the experimentally identified RNA-binding peptides (referred to as RBDpeps) studies ^6,44–46^ were annotated as high-confidence RBDs. Similarly, high-confidence non-RNA-binding domains were those homology-annotated domains in the RBPs that are not in the RNA-binding and -processing domain list and not supported by RBDpeps studies. Within the occlusion map output, we observed the percentage of the high-confidence RBDs that have at least one pRBR (p-value < 0.05) or stringent pRBRs (p-value < 0.001) (akin to sensitivity), and also the percentage of the high-confidence non-RBDs have at least one pRBR or stringent pRBRs (akin to false positive rate (FPR)). Positive likelihood ratio (also called fold-enrichment) was then calculated as the ratio of sensitivity versus FPR, to demonstrate how well the pRBRs are enriched in high-confidence RBDs over high-confidence non-RBDs in the proteins. Similarly, we ran the baseline tools RNABindPlus and DRNApred, which are designed to predict RNA-binding residues in RBPs, on the same set of RBPs. We obtained positive likelihood ratio by observing the percentage of the high-confidence RBDs that have at least one predicted RNA-binding residue (i.e. sensitivity) and the percentage of the high-confidence non-RBDs that have at least one predicted RNA-binding residue (i.e. FPR). By using different p-value/ score threshold when defining pRBR and predicted RNA-binding residues, we got the positive likelihood ratio changes at different sensitivity level.

### eCLIP Data Processing and Analysis

eCLIP sequence reads were processed and analyzed as previously described ^48^. For each protein of YWHAG/H/E/Z and HSP90AA1, a total of six eCLIP libraries, 2 replicates for each of the IP, SMInput and control group, were sequenced, while YWHA family shares the same control libraries. For each protein of INO80B, PIAS4, NR5A1, ACTN3, and MCCC1, 2 or 3 replicates for each of the IP, SMInput were sequenced, Reads were first mapped to repetitive elements and only the unmapped reads were retained and were next mapped to human genome (hg19). The uniquely mapped reads were kept for downstream analysis. PCR duplicates were further removed to obtain ‘usable reads’ with use of a randomer (N10) sequence positioned one of the adapter oligos. Next, usable reads were counted for both eCLIP (immunoprecipitation) and size-matched input (SMInput) samples across different genomic regions (i.e. 5′UTR, 3′UTR, coding exons (CDS), and intronic regions) for all coding genes annotated in UCSC Known Gene table (hg19). To identify the enrichment of binding signals in CLIP samples above SMInput in certain genomic region of certain gene, fold-enrichment was calculated as the ratio of read counts within this region in CLIP versus SMInput. Only regions with at least 10 reads in both of eCLIP and SMInput samples were considered. Sequencing read peaks were identified using CLIPper algorithm ^78^ for both eCLIP and SMInput libraries. CLIPper-defined peaks were then normalized to size-matched input (SMInput) by comparing the read density in immunoprecipitation (IP) and SMInput samples (referred to as INPUT normalization). Significant peaks were defined as the peaks whose number of reads from the IP sample were 8-fold greater than the number of reads from the SMInput sample with a p-value < 0.001. P-value was calculated by Yates’ Chi-Square test (Fisher Exact Test was performed when the observed or expected read number was below five). For HSP90AA1 and YWHA G/H/E/Z, reproducible peak across both biological replicates were identified using an irreproducible discovery rate (IDR) approach used in our previous study ^79^. To validate the new RBPs (i.e. INO80B, PIAS4, NR5A1, ACTN3, and MCCC1) and their predicted functional domains, we applied a more stringent strategy to identify the reliable reproducible peaks using eCLIP output from size-matched “V5 tag only” experiments (shown in **Figure S9C**). In this strategy, the sequencing reads are processed in the same way above except we did two sets of normalization in the INPUT normalization step: (1) the standard normalization, i.e. protein IP against protein SMinput; (2) the normalization against V5 tag only output, i.e. protein IP against the IP sample from the size-matched V5 tag experiment. ChIP-R^49^ algorithm was then used to identify the reproducible peaks for each of these outputs because of its capability of handling more than 2 replicates. The shared reproducible peaks from these two normalization ways form the final list of reproducible peaks where at least 5 mapped reads are found in the peak for each IP sample of the protein are also required.

### Peak Enrichment Analysis

Significantly enriched eCLIP peaks (described above) were assigned to different transcriptomic regions (i.e. 5′UTR, 3′UTR, coding exons (CDS), and intronic regions) with BEDTools ^80^ using annotations from GENCODE. For peaks whose coordinates overlap with multiple transcriptomic regions, the following priority were applied to assign these peaks each with a single region: CDS, 5′UTR, 3′UTR, then proximal (as defined as less than 500 bp from an exon–intron boundary) or distal introns (as defined as 500 bp or greater from an exon– intron boundary). The fraction of peaks in a specific transcriptomic region (*P_CLIP_*) was obtained by dividing the number of peaks in this region by the total number of peaks in all regions (5′UTR, 3′UTR, CDS, proximal intron and distal intron). Fold-enrichment (*E*) was then calculated as *E_region_* = *log*_2_(*P_region_*/*S_region_*) to measure the binding preference of the tested RBP over this transcriptomic region, where *S_region_* is the fractional region size derived by dividing the total number of base pairs in that region relative to the total number of base pairs in all regions. *E_region_* with positive values indicate overrepresentation of peaks in this region and negative values indicate the underrepresentation. To be noticed, if a base pair is associated with multiple transcriptomic regions, this base pair was assigned to regions in the following priority: CDS, 5′UTR, 3′UTR, then proximal or distal introns.

### GO enrichment for candidate RBPs

We retrieved the enriched Gene Ontology (GO) terms for the candidate RBPs (YWHA family, INO80B, PIAS4, NR5A1, ACTN3, and MCCC1) with overrepresentation test implemented by PANTHER classification system ^81^. To simplify the output, we only observed the enrichment in GO terms from level 5 of the GO hierarchy system.

Gene ontology enrichment analysis for HSP90A was performed using the Enrichr Gene Ontology enrichment tool (Kuleshov et al., 2016). Results were ranked by the “combined score”, which combines p-value and z-score by multiplication, i.e. combined score = log(p) * z-score.

### Other Statistical Analysis

Other Statistical analysis was performed using Scipy package in Python 3.

## Supplementary Tables

**Table S1.** Classification performance of each components of HydRA.

**Table S2.** HydRA Classification scores for human proteins.

**Table S3.** Statistics of occlusion scores for human domains.

**Table S4.** Feature and model selection for structure-based GNN models.

## KEY RESOURCES TABLE

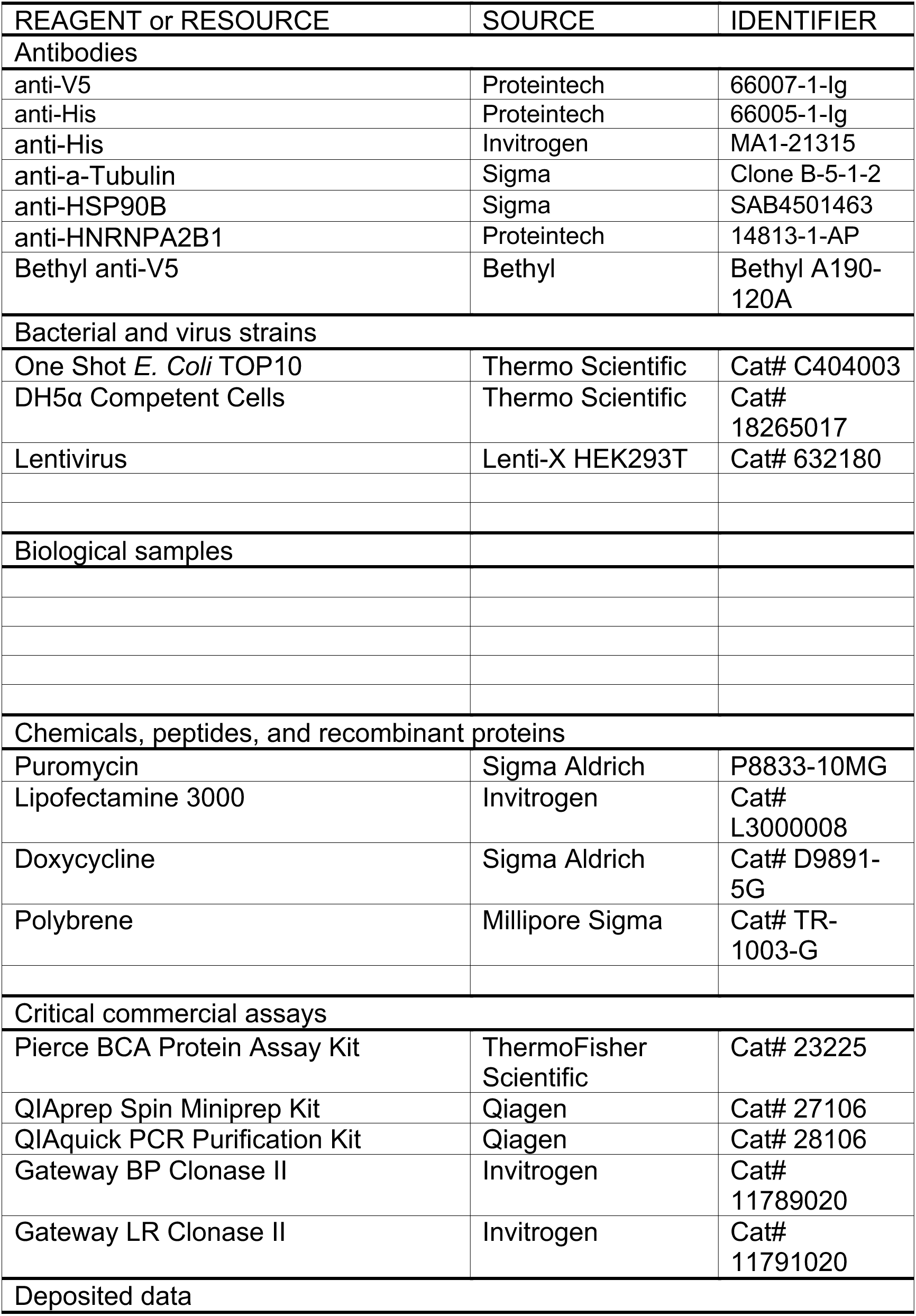
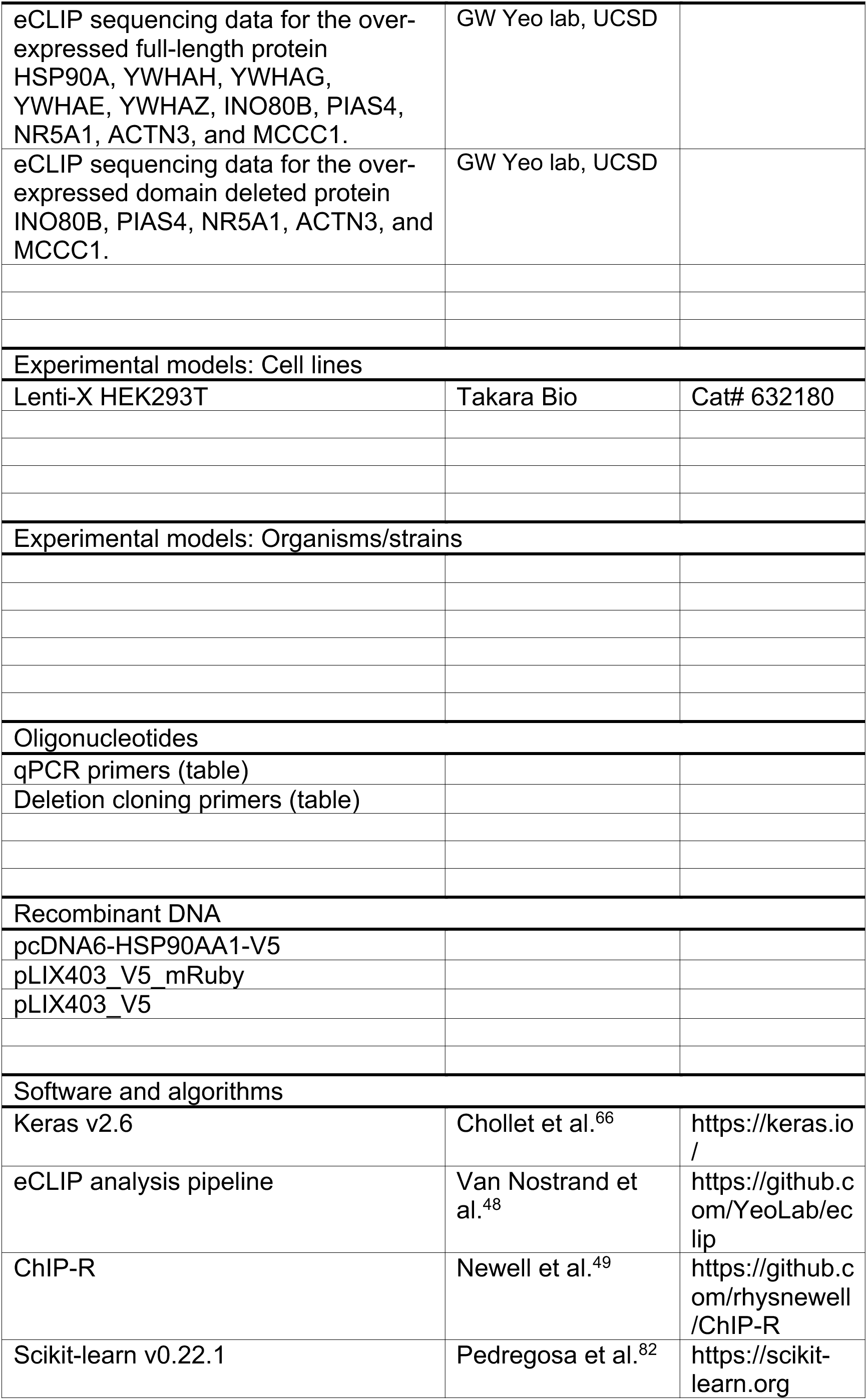
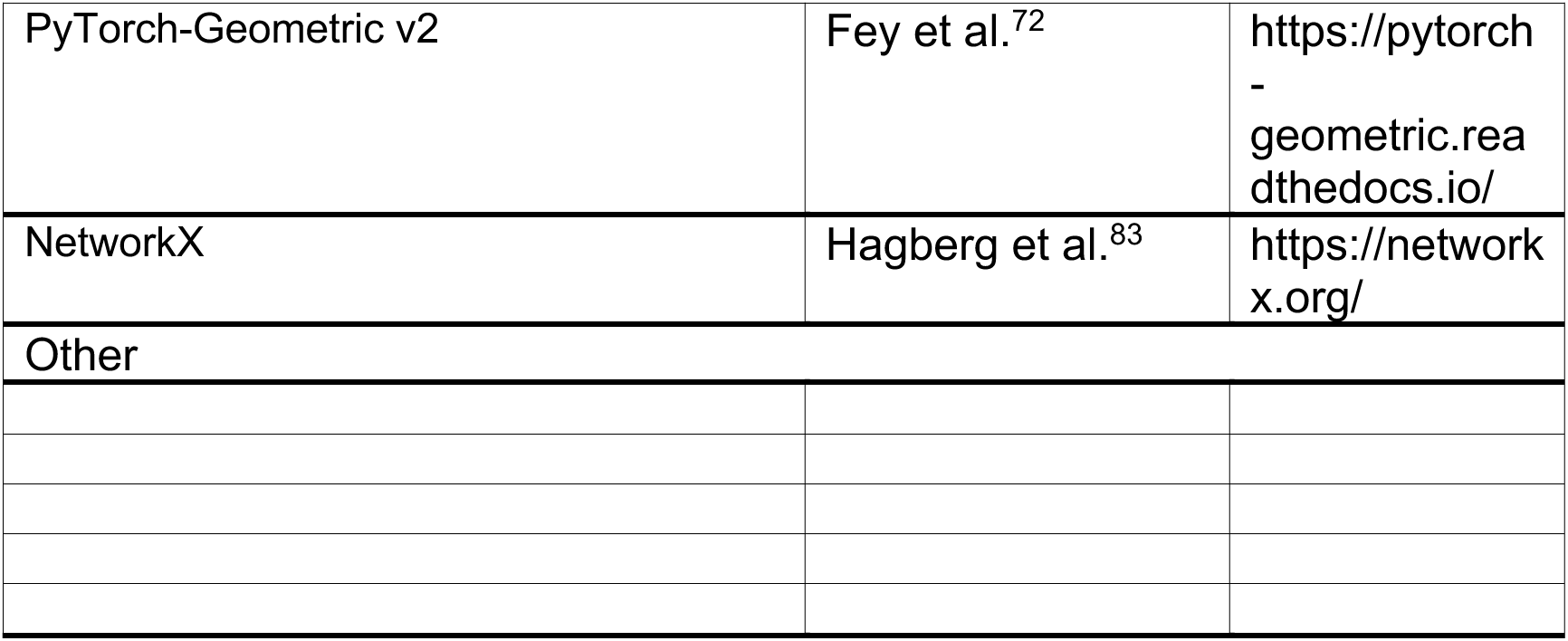

